# Apoptosis, G1 phase stall and premature differentiation account for low chimeric competence of Human and rhesus monkey naïve pluripotent stem cells

**DOI:** 10.1101/2020.03.27.011890

**Authors:** Irène Aksoy, Cloé Rognard, Anaïs Moulin, Guillaume Marcy, Etienne Masfaraud, Florence Wianny, Véronique Cortay, Angèle Bellemin-Ménard, Nathalie Doerflinger, Manon Dirheimer, Chloé Mayère, Pierre-Yves Bourillot, Cian Lynch, Olivier Raineteau, Thierry Joly, Colette Dehay, Manuel Serrano, Marielle Afanassieff, Pierre Savatier

**Author notes:** **Corresponding authors:** Irène Aksoy and Pierre Savatier, Stem Cell and Brain Institute, INSERM U1208, 18 Avenue du Doyen Lépine, F-69675 Bron Cedex, France. Material requests should be addressed to Irène Aksoy.

## Abstract

After reprogramming to naïve pluripotency, human pluripotent stem cells (PSCs) still exhibit very low ability to make interspecies chimeras. Whether this is because they are inherently devoid of the attributes of chimeric competency or because naïve PSCs cannot colonize embryos from distant species remains to be elucidated. Here, we have used different types of mouse, human and rhesus monkey naïve PSCs and analyzed their ability to colonize rabbit and cynomolgus monkey embryos. Mouse embryonic stem cells (ESCs) remained mitotically active and efficiently colonized host embryos. In contrast, primate naïve PSCs colonized host embryos with much lower efficiency. Unlike mouse ESCs, they slowed DNA replication after dissociation and, after injection into host embryos, they stalled in the G1 phase and differentiated prematurely, regardless of host species. We conclude that human and non-human primate naïve PSCs do not efficiently make chimeras because they are inherently unfit to remain mitotically active during colonization.

## Introduction

Human embryo-derived pluripotent stem cells (PSCs) and human induced PSCs (iPSCs) exhibit biological and functional characteristics of primed pluripotency: 1) dependence on fibroblast growth factor 2 (FGF2)/extracellular signal-regulated kinase and activin A/SMAD signaling for self-renewal; 2) inactivation of the 2nd X chromosome in female lines; and 3) a global transcriptome more similar to that of post-implantation epiblast in the gastrulation embryo (Chen and Lai, 2014; Davidson et al., 2015; Nakamura et al., 2016; Nichols and Smith, 2009). Although PSC lines obtained from rhesus monkeys are less well characterized, they also exhibit the essential characteristics of primed pluripotency (Wianny et al., 2008). In this respect, human and non-human primate PSCs differ from their murine counterparts, which exhibit biological and functional characteristics of naïve pluripotency (Chen and Lai, 2014; Davidson et al., 2015; Nichols and Smith, 2009). Notably, naïve and primed PSCs differ in their capacity to generate chimeras following injection into pre-implantation embryos. Although mouse ESCs (mESCs) produce germline chimeras, rhesus monkey PSCs do not colonize the ICM/epiblast of the host embryo and undergo apoptosis (Tachibana et al., 2012).

Different culture media with capacity to produce primed-to-naïve conversion of human, cynomolgus, and rhesus macaque PSCs have been reported. The media are variously termed naïve human stem cell media (NHSM) (Gafni et al., 2013), E-NHSM (https://hannalabweb.weizmann.ac.il), NHSM-v (Chen et al., 2015b), 3iL (Chan et al., 2013), Reset (Takashima et al., 2014), 5i/L/A, and 6i/L/A (Theunissen et al., 2016; Theunissen et al., 2014), 4i/L/b (Fang et al., 2014), TL2i (Chen et al., 2015a), 2iLD, 4i and FAC (Wu et al., 2017), t2iLGöY (Guo et al., 2017), and LCDM (Yang et al., 2017). PSCs under these culture conditions display characteristic features of naïve state pluripotency of mESCs with a reconfigured transcriptome and epigenome (Chen et al., 2015a; Huang et al., 2014; Nakamura et al., 2016), loss of FGF2 dependency (Chen et al., 2015a; Takashima et al., 2014), gain of STAT3 dependency (Chan et al., 2013; Chen et al., 2015a; Gafni et al., 2013; Takashima et al., 2014), reactivation of the second X-chromosome (Fang et al., 2014; Takashima et al., 2014; Theunissen et al., 2014), and elevation of oxidative phosphorylation (Takashima et al., 2014; Ware et al., 2009).

Colonization of mouse embryos by human naïve PSCs and their subsequent participation in germ layer differentiation has been reported (Fang et al., 2014; Gafni et al., 2013). However, these early results were not confirmed by subsequent reports: low rates of chimerism (<0.001%) in <1% of the fetuses were reported after injection of human naïve PSCs in mouse blastocysts (Masaki et al., 2015; Theunissen et al., 2016). Similarly, very low rates of chimerism were also observed in pig fetuses after injecting human naïve iPSCs in pig embryos (Wu et al., 2017). Whether human PSCs have failed to produce chimeras because naïve PSCs in general cannot colonize embryos from distant species or because human PSC are inherently devoid of the attributes of chimeric competence is not known.

To explore this issue, we investigated the differential ability of mESCs, rhesus monkey, and human naïve PSCs to colonize both closely and distantly related host embryos. To circumvent the ban on introducing human embryo-derived PSCs into animal embryos, we used embryo-derived rhesus monkey PSCs and human iPSCs. As a reference, we explored the ability of mESCs, the gold standard of naïve pluripotency, to colonize evolutionary distant host embryos. As host species, we selected rabbit and cynomolgus monkey embryos as hosts. Primates and glires (rodents + lagomorphs) diverged between 85 and 97 million years, while rodents and lagomorphs diverged between 77 and 88 million years (www.timetree.org). The divergence time makes rabbit embryos an almost equidistant environment for comparing the colonizing capabilities of rodent and primate PSCs. In addition, rabbits have several advantages in testing the colonization capacity of human PSCs. The early rabbit embryos share common features with primate embryos in their developmental characteristics (Madeja et al., 2019); unlike the three-dimensional egg-cylinder shape of rodent embryos during gastrulation, primate and rabbit embryos develop into a flattened disc at the surface of the conceptus; primate and rabbit embryos are also markedly similar with respect to the timing of zygotic genome activation and the timing and regulation of X-chromosome inactivation. In rodents and primates, gastrulation occurs in the implanted embryo buried within the uterine wall. However, in rabbits, gastrulation begins shortly before implantation, allowing easier access to a wider developmental window. Furthermore, the rabbit PSC lines derived from pre-implantation embryos require FGF2 and TGFβ for inhibition of differentiation and exhibit the cardinal features of primed pluripotency (Osteil et al., 2016; Osteil et al., 2013). Thus, rabbit pre-implantation embryos appear to be more similar to primate embryos than mouse embryos with respect to the mechanisms regulating pluripotency. Therefore, they may be a better host for examining the colonization competence of human PSCs. As a closely related host for Human and rhesus monkey PSCs, we used cynomolgus macaque embryos. The common ancestor of macaques and Humans dates back 29 million years, while rhesus and cynomolgus macaques diverged 3.7 million year ago (www.timetree.org). We explored the differential ability of mESCs, rhesus monkey PSCs and human naïve iPSCs to colonize rabbit and cynomolgus embryos, and drew conclusions on the inherent abilities of rodent and primate naïve PSCs to colonize distantly-related hosts.

## Results

### Colonization of rabbit embryos by mESCs

Fresh rabbit embryos observed by epifluorescence microscopy typically show a high level of autofluorescence (**Supplemental data S1A**), which could be confounding for the detection of GFP-expressing cells in chimeras. To overcome this limitation, we systematically used an anti-GFP antibody. No immunostaining was observed when applied to rabbit embryos (i) prior to any injection of cells, and (ii) after injection of 10 rhesus wild-type PSCs (LyonES (Wianny et al., 2008)) that do not express any fluorescent reporters. Immunolabeling was observed in both rhesus PSCs expressing GFP as a fusion between tau protein and GFP (LyonES-tauGFP) **(Supplemental data S1B)** and rabbit embryos colonized with naïve-like rabbit iPSCs expressing GFP **(Supplemental data S1C)**(Osteil et al., 2016). Thus, immunolabeling of GFP allows an effective detection of GFP expression in our system.

Mouse ESCs grown in serum/LIF and, specially, in 2i/LIF epitomize naïve pluripotency and, for this reason, we tested their ability to colonize rabbit embryos. For this purpose, 10 mESC-GFP cells cultured in serum/LIF condition were injected into rabbit embryos at the morula stage (E2). The morulas were subsequently cultured for 1–3 days *in vitro* (DIV) and developed into early (E3), mid (E4), and late (E5) blastocysts (1, 2, and 3 DIV, respectively). The vast majority of rabbit blastocysts had incorporated groups of GFP^+^ cells (97.5%, n = 81; **Fig. 1A**, **Table S1**). mESCs and their progeny divided very actively, as shown by both the expansion of the GFP^+^ cell pool during *in vitro* culture of chimeric embryos and a high incorporation of 5-ethylnyl-2ʹ-deoxyuridine (EdU) in most GFP^+^ cells (**Figs. 1B,C**; **Supplemental data S2**). It should be noted that, under the conditions used (very short incorporation time), the EdU labeling of host embryo cells is markedly low. GFP^+^ cells and their progeny expressed the transcription factors and pluripotency markers Oct4, Nanog and SRY-box transcription factor (Sox)2, as revealed by immunostaining at E5 (3 DIV) (**Figs. 1D, E**). In contrast, GFP^+^ cells did not express Sox17, a primitive endoderm marker, indicating that they did not differentiate to the primitive endoderm lineage. We performed a similar experiment using cynomolgus monkey embryos as hosts. For this purpose, 10 mESC-GFP cells were injected into cynomolgus embryos at the morula stage (E4). The morulas were subsequently cultured for 3 DIV and developed into mid/late (E7) blastocysts before immunostaining with GFP, OCT4, NANOG, SOX2, and SOX17 antibodies. Among the five embryos analyzed, three contained 8–16 GFP^+^ cells in the ICM. The first embryo was immunostained with GFP, OCT4, and SOX2 antibodies and contained 16 GFP^+^/OCT4^+^/SOX2^+^ cells. The second one was immunostained with GFP and NANOG antibodies and contained eight GFP^+^/NANOG^+^ cells. The third one was immunostained with GFP and SOX17 antibodies and contained eight GFP^+^/ SOX17^−^ cells (**Fig. 1F**, **Table S1**). In another experiment, 10 mESC-GFP cells propagated in N2B27 supplemented with LIF, PD0325901, and CHIR99021 (*i.e.*, 2i/LIF condition) were injected into rabbit morulas and subsequently cultured for 3 DIV prior to immunostaining (**Supplemental data S3A**, **Table S1**). Of the 19 E5 blastocysts obtained, 14 had incorporated groups of GFP^+^ cells. Immunostaining showed that 100% of chimeric embryos harbored GFP^+^ cells expressing SOX2 (n = 7/7), whereas none of them expressed SOX17 (n = 0/7). Overall, these results show that mESCs, whether in serum/LIF or 2i/LIF conditions, continue to express pluripotency markers and expand after injection into evolutionary distant embryos, whether rabbit or cynomolgus monkey embryos.

**Figure 1:**
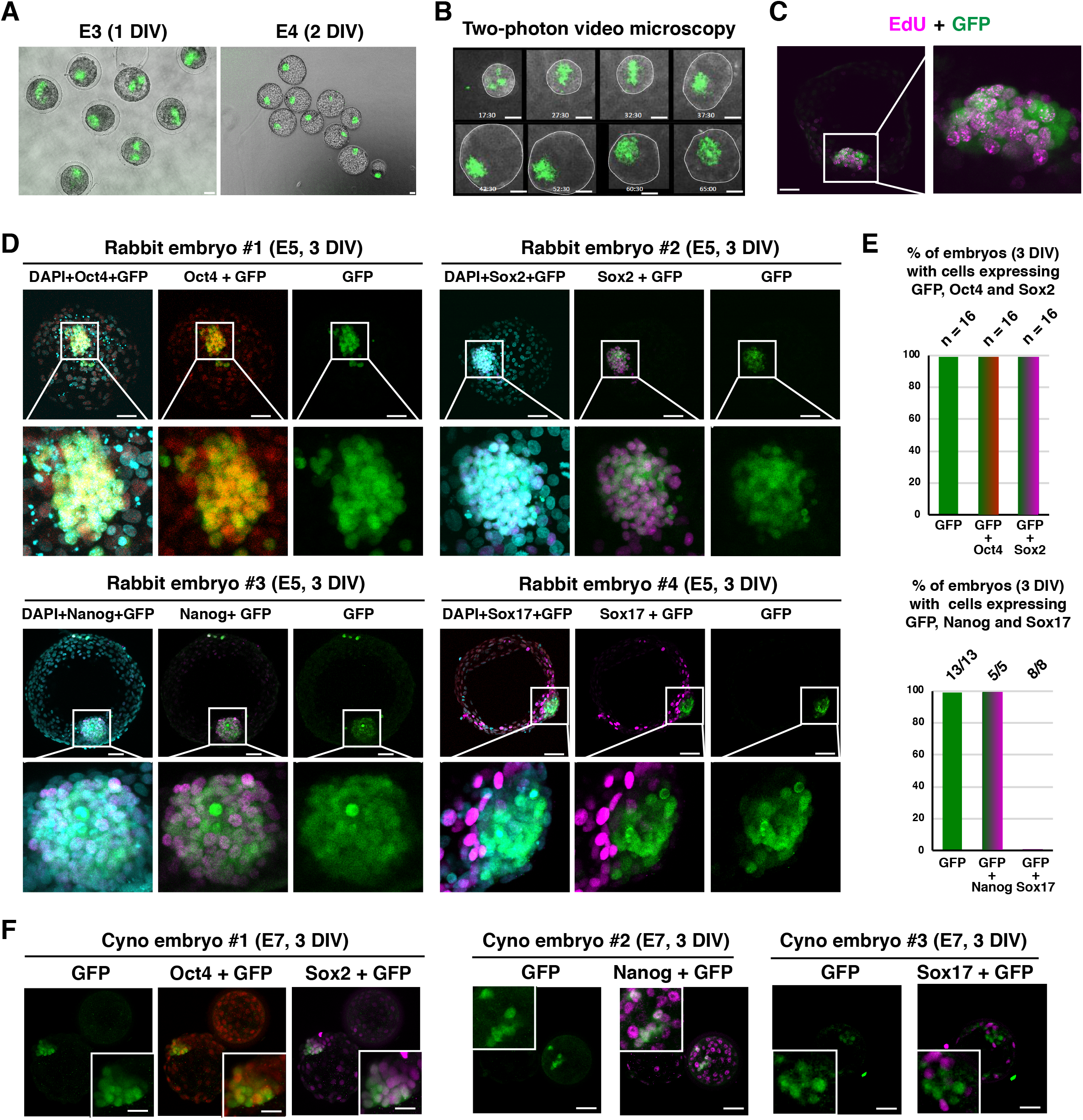
Colonization of rabbit and cynomolgus embryos by mouse ESCs (mESCs). (**A**) Epifluorescence images of the early-(E3, 1 DIV) and mid-blastocyst-stage rabbit embryos (E4, 2 DIV) resulting from microinjection of 10 mESC-GFP cells (scale bars: 50 µm). (**B**) Two-photon microscope images of the late blastocyst-stage rabbit embryos (E2–E5, 0–3 DIV) resulting from microinjection of 10 mESC-GFP cells (scale bar: 50 µm). (**C**) Immunostaining of GFP and EdU in a late blastocyst-stage rabbit embryo (E5, 3 DIV) after microinjection of 10 mESC-GFP cells into the morula-stage (E2) embryo (confocal imaging; scale bars: 50 μm). (**D**) Immunostaining of GFP, Oct4, Sox2, Nanog, and Sox17 of late blastocyst-stage rabbit embryos (E5, 3 DIV) after microinjection of 10 mESC-GFP cells into morula-stage (E2) embryos (confocal imaging; scale bars: 50 μm) (n = 135). (**E**) Histogram of percentage of rabbit embryos with GFP^+^/Oct4^+^, GFP^+^/Sox2^+^, GFP^+^/Nanog^+^, and GFP^+^/Sox17^+^ cells at 3 DIV (n = 74). (**F**) Immunostaining of GFP, Oct4, Sox2, and Sox17 of late blastocyst-stage (E7) cynomolgus embryos after microinjection of 10 mESC-GFP cells into morula-stage (E4) embryos (confocal imaging; scale bars: 50 μm) (n = 7).

We subsequently studied the ability of mESCs to colonize the embryonic disk of rabbit pregastrula embryos (E6) and the three germ layers after gastrulation (E9). For this purpose, the E2 embryos injected with 10 mESC-GFP cells (grown in serum/LIF) were transferred into the oviducts of surrogate rabbits. Nine embryos were recovered after 4 days (*i.e.*, E6). All contained GFP^+^ cells expressing OCT4 and SOX2 (**Supplemental data S3B**, **Table S1**). Three embryos were immunostained with a SOX17 antibody and did not show SOX17^+^ cells. On the other hand, the GFP^+^ cells formed a coherent group of cells in the embryonic disk, suggesting that mouse cells did not mix with rabbit cells. To assess whether mESCs participated in the development of rabbit embryos after gastrulation, 20 fetuses were recovered 7 days after transfer to surrogate mothers (*i.e.*, E9) (**Supplemental data S3C**, **Table S1**). Fetuses contained GFP^+^/SOX2^+^, GFP^+^/TUJ1^+^ and GFP^+^/NANOG^−^ cells in the neurectoderm (n = 12/12). None of the fetuses had GFP^+^ cells in embryonic tissues derived from the mesoderm and endoderm or in extra-embryonic tissues. These results strongly suggest that mESCs injected into rabbit morulas contribute to the expansion of the late epiblasts until the onset of gastrulation. After gastrulation, mESCs were able to contribute to the neuroectoderm, but not to other embryonic and extra-embryonic lineages.

### Reprogramming rhesus PSCs to naïve pluripotency

The LyonES rhesus line expressing a tauGFP transgene under the control of the ubiquitous CAG promoter has been previously described (Wianny et al., 2008). LyonES-tauGFP cells were infected with the pGAE-STAT3-ER^T2^ lentiviral vector (Chen et al., 2015a), and a clonal cell line stably expressing STAT3-ER^T2^ was isolated as LyonES-tGFP-(S3). Immunostaining with anti-STAT3 antibody showed increased labeling and nuclear translocation following the treatment of LyonES-tGFP-(S3) cells with tamoxifen (**Supplemental data S4A**). LyonES-tGFP-(S3) cells were cultured in the presence of tamoxifen to activate STAT3-ER^T2^ and with 1,000 U/mL of LIF, MEK, and GSK3β inhibitors. Consistent with previous observations in human PSCs (Chen et al., 2015a), the LyonES-tGFP-(S3) cells formed small dome-shaped colonies, which were termed rhesus TL2i (rhTL2i) cells (**Fig. 2A**). These colonies demonstrated identical morphology with that previously described for human TL2i cells (Chen et al., 2015a). We developed a variant of the TL2i protocol, in which MEKi and GSK3βi were replaced by CNIO-47799, a chemical inhibitor of cyclin-dependent kinase (CDK)8 and CDK19 (CDK8/19i) for reprogramming to the naïve state (Lynch et al., In revision). This protocol is referred to as TL−CDK8/19i and the reprogrammed cells as rhesus TL-CDK8/19i (rhTL-CDK8/19i). LyonES-tGFP-(S3) cells were submitted to six other reprogramming protocols previously developed to revert human or monkey PSCs to naïve pluripotency, including NHSM-v and E-NHSM (Chen et al., 2015b; Gafni et al., 2013), 4i/L/b (Fang et al., 2014), t2iLGöY (Guo et al., 2016; Takashima et al., 2014), 5iL/A or 6iL/A (Theunissen et al., 2014), and LCDM (extended pluripotent stem [EPS]) (Yang et al., 2017). It should be noted that the six protocols were applied to LyonES-tGFP-S3 cells in the absence of tamoxifen, thus maintaining STAT3-ER^T2^ in an inactive state. LyonES-tGFP-S3 cells differentiated or underwent apoptosis when cultured in LIF/MEKi/GSK3βi (not shown). Of the eight protocols tested, seven (*i.e.,* NHSM-v, E-NHSM, 4i/L/b, t2iLGöY, LCDM [EPS], TL2i, and TL-CDK8/19i) produced rhesus PSCs forming dome-shaped colonies (**Fig. 2A**). Continued instability was observed with protocol 5iL/A, which was subsequently omitted from the study.

**Figure 2:**
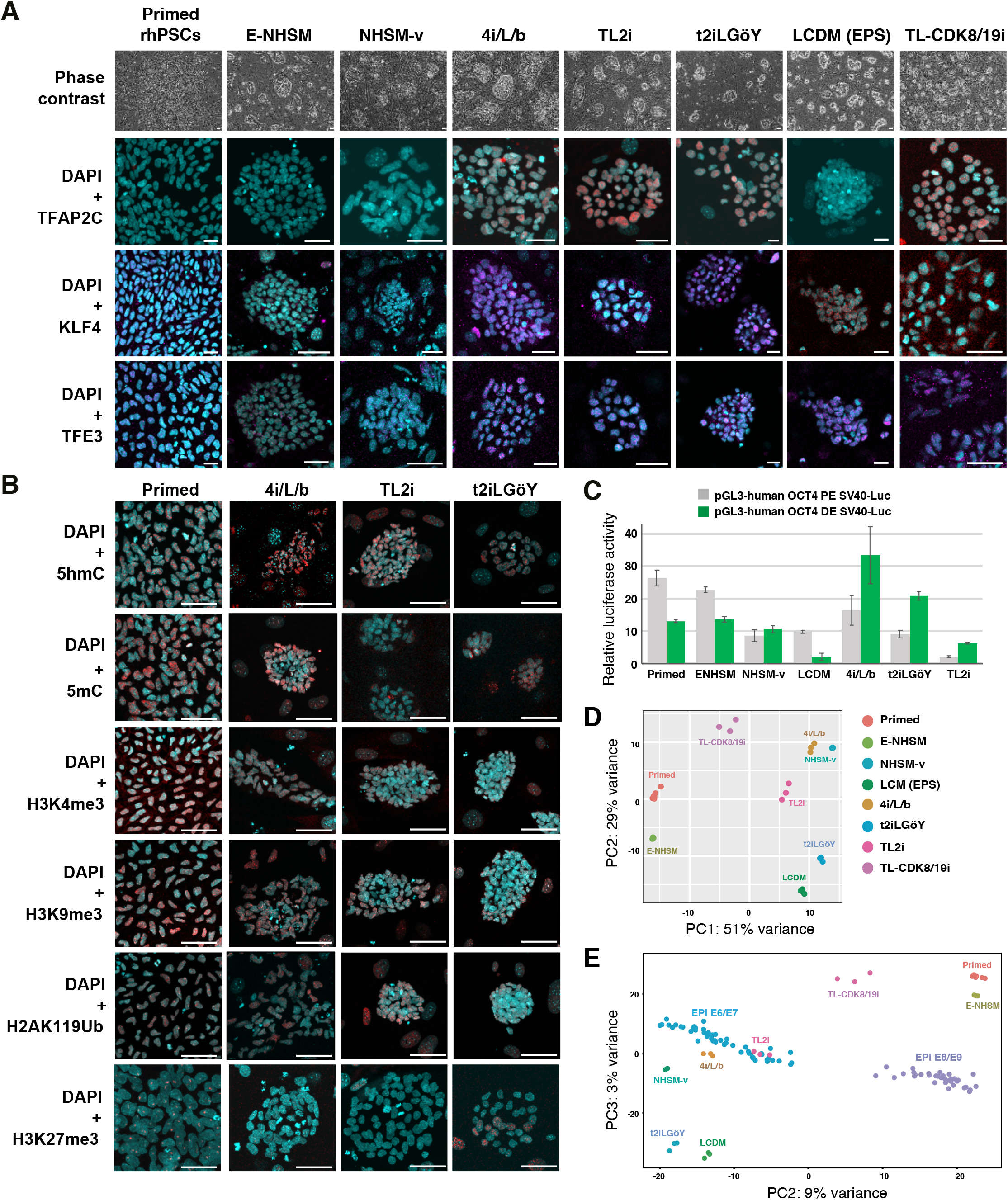
Characterization of rhesus PSCs after reprogramming to naïve pluripotency. (**A**) Phase contrast and immunostaining of naïve pluripotency markers, TFAP2C, KLF4, and TFE3 in LyonES-tGFP-(S3) cells, before (primed rhPSCs) and after reprogramming to the naïve state (E-NHSM, NHSM-v, 4i/L/b, TL2i, t2iLGöY, LCDM (EPS), and TL-CDK8/19i) (confocal imaging; scale bars: 50 μm). (**B**) immunostaining of 5ʹ-methylcytosine (5mC), 5ʹ-hydroxymethylcytosine (5hmC), H3K4me3, H3K9me3, H2AK119Ub, and H3K27me3 in LyonES-tGFP-(S3) cells, before (primed) and after reprogramming to the naïve state (4i/L/b, TL2i, and t2iLGöY) (confocal imaging; scale bars: 50 μm). (**C**) Activity of the proximal and distal enhancer (PE and DE, respectively) of *OCT4* measured in a luciferase assay after transient transfection in primed PSCs, E-NHSM, NHSM-v, 4i/L/b, TL2i, t2iLGöY, and LCDM cells. (**D**) Principal component analysis (PCA) for primed PSCs, E-NHSM, NHSM-v, 4i/L/b, TL2i, t2iLGöY, LCDM, and TL-CDK8/19i populations based on RNA-sequencing data. (**E**) PCA for primed PSCs, E-NHSM, NHSM-v, 4i/L/b, TL2i, t2iLGöY, LCDM, and TL-CDK8/19i populations based on bulk RNA-seq and single-cell RNA-seq data of cynomolgus epiblast from E6 to E9 embryos (Nakamura et al., 2016).

Unlike conventional rhesus PSCs, which were mechanically passaged in clumps after collagenase dissociation, all reprogrammed cells were routinely dissociated into single cells using trypsin or trypLE. The colonies exhibited variable expression of naïve transcription factors and cell surface molecules (**Fig. 2A and Supplemental data S4B, G**). Notably, the transcription factors AP-2 gamma (TFAP2C), KLF4, and transcription factor binding to IGHM enhancer 3 (TFE3) displayed stronger expression after reprogramming using the 4i/L/b, t2iLGöY, TL2i, and TL-CDK8/19i protocols. LCDM (EPS) cells expressed KLF4 and TFE3 but not TFAP2C. Cells reprogrammed with the other protocols did not express any of these naïve markers (TFAP2C, KLF4, TFE3). t2iLGöY, TL2i, and 4i/L/b cells featured lower levels of H3K4me3 marks (**Fig. 2B, Supplemental data S4C–D**). They also displayed diffuse H3K27me3 immunostaining, compared with punctate staining in primed, NHSM-v, and ENHSM cells (**Fig. 2B, Supplemental data S4C, E**). Immunostaining of 5ʹ-methylcytosine and 5ʹ-hydroxymethylcytosine also revealed global genome hypomethylation in t2iLGöY, LCDM (EPS), and TL2i cells (**Fig. 2B, Supplemental data S4C–D**). Primed, ENHSM, and NHSM-v cells exhibited punctate immunostaining for H2AK119Ub, indicating the presence of an inactive X chromosome, whereas t2iLGöY, LCDM (EPS), TL2i, and 4i/L/b cells displayed diffuse immunostaining, suggesting X chromosome reactivation (**Fig. 2B, Supplemental data S4C, E**). Altogether, these results strongly suggest that t2iLGöY, LCDM (EPS), TL2i, and 4i/L/b cells underwent epigenome reconfiguration, in line with the results of naïve marker expression. Finally, consistent with their naïve status, t2iLGöY, TL2i, and 4i/L/b cells exhibited preferential usage of the distal enhancer of Oct4 in a luciferase assay **(Fig. 2C).**

To further characterize reprogramming, gene expression was analyzed by bulk RNA sequencing to assess transcriptome reconfiguration. Principal component analysis (PCA) and hierarchical clustering showed that primed and E-NHSM cells clustered on one side, whereas NHSM-v, 4i/L/b, t2iLGöY, TL2i, and LCDM (EPS) cells clustered on the other side (**Fig. 2D**, **Supplemental data S4F, Table S2**). TL-CDK8/19i cells demonstrated an intermediate position, suggesting that transcriptome reconfiguration was less pronounced. A comparative analysis was performed between the primed and naïve rhesus PSC lines and single-cell RNA-sequencing data of the cynomolgus epiblast (Nakamura et al., 2016). The NHSM-v, 4i/L/b, t2iLGöY, TL2i, and LCDM (EPS) cells also clustered closer to the epiblast cells of the E6/E7 cynomolgus embryo (Nakamura et al., 2016) than the E-NHSM and primed cells (**Fig. 2E**). These results show that, consistent with the immunostaining data, reprogramming of rhesus PSCs is characterized by a transcriptome shift reflecting the acquisition of characteristic features of the pre-implantation epiblast as previously reported with human naïve PSCs (Nakamura et al., 2016).

### Premature differentiation of rhesus naïve PSCs after injection into rabbit embryos

The ability of reprogrammed rhesus PSCs to colonize rabbit ICM/epiblast was tested. For this purpose, 10 LyonES-tGFP-(S3) cells of each type [primed, NHSM-v, E-NHSM, LCDM (EPS), 4i/L/b, t2iLGöY, TL2i, and TL-CDK8/19i] were injected into rabbit morulas (E2). The embryos were subsequently cultured for 1–3 DIV in gradual media and developed into early (E3), mid (E4), and late blastocysts (E5). There was no difference observed in the rate of development between control (uninjected) and injected embryos (not shown). In the first set of experiments, the fate of injected cells was monitored by two-photon microscopy for 48–72 h. Contrary to our observations with mESCs, rhesus PSCs did not actively proliferate in host embryos, and most of them did not incorporate into ICM/epiblast (**Supplemental data S5– S8**). GFP^+^ embryos and cell counts confirmed this finding: the number of embryos with GFP^+^ cells decreased with time in culture, ranging from 0 (E-NHSM) to 62% (t2iLGöY) at 3 DIV compared with 22–96% at 1 DIV (**Fig. 3A**, **Table S1**). In addition, the percentage of embryos with ≥4 GFP^+^ cells diminished in all experimental conditions tested, ranging from 0 (E-NHSM) to 24% (TL2i) at 3 DIV *vs.* 0 (E-NHSM) to 79% (t2iLGöY) at 1 DIV (**Fig. 3B**). These results indicate that rhesus PSCs failed to actively divide and/or were gradually eliminated during embryonic development, regardless of the reprogramming strategy used for conversion to the naïve state. Despite the failure of the rhesus PSCs to thrive in a rabbit embryo environment, variations in the survival rate of GFP^+^ cell at 3 DIV were observed between reprogramming strategies. There were no GFP^+^ cells observed in embryos injected with either primed rhesus PSCs (0/90) or E-NHSM cells (0/71). In contrast, GFP^+^ cells were observed at varying frequencies using other reprogramming protocols. At 3 DIV, the highest rates were observed with TL2i (57%, n = 490), 4i/L/b (41%, n = 228), t2iLGöY (37%, n = 168), LCDM (26%, n = 335), and TL-CDK8/19i (20%, n = 260). TL2i was the only reprogramming method that returned more GFP^+^ cells at 2 DIV (58%, n = 145) and 3 DIV (57%, n = 204) compared with 1 DIV (44%, n = 141). These results suggested that TL2i cells have a slightly increased proliferative capacity compared with cells reprogrammed using other methods (**Fig. 3B**).

**Figure 3:**
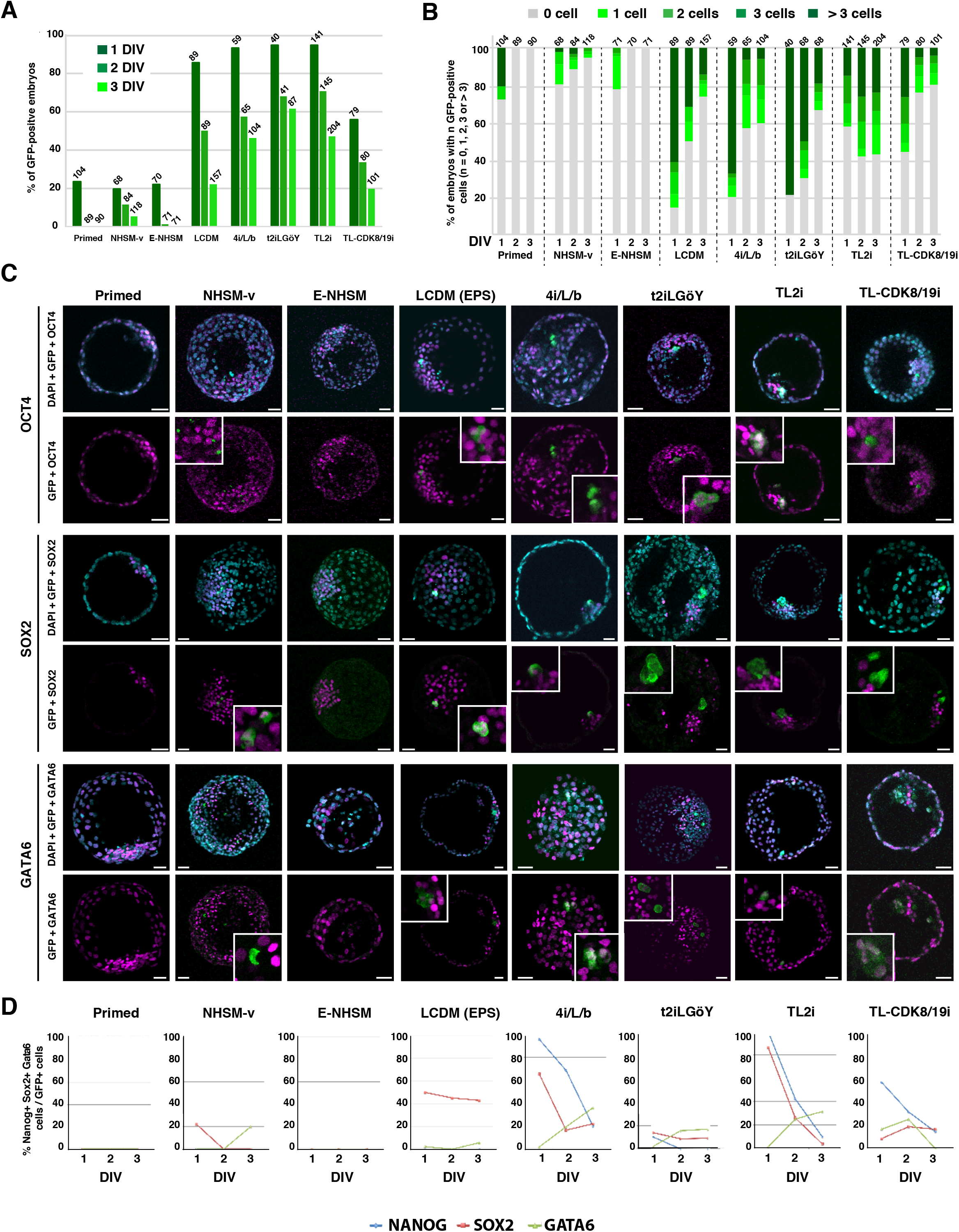
Colonization of rabbit embryos by rhesus naïve PSCs. (**A**) Percentage of rabbit embryos with GFP^+^ cells, 1–3 days (1–3 DIV) after injection of rhesus LyonES-tGFP-(S3) cells into morula-stage (E2) embryos, before (Primed) and after reprogramming to the naïve state (E-NHSM, NHSM-v, 4i/L/b, TL2i, t2iLGöY, LCDM (EPS), and TL-CDK8/19i) (n = 650 at 1 DIV; n = 664 at 2 DIV; n = 932 at 3 DIV). (**B**) Percentage of rabbit embryos with 0, 1, 2, 3, or >3 GFP^+^ cells 1–3 days (1–3 DIV) after injection of rhesus LyonES-tGFP-(S3) cells into morula-stage (E2) embryos (n = 650 at 1 DIV; n = 664 at 2 DIV; n = 932 at 3 DIV). (**C**) Immunostaining of GFP, Oct4, Sox2, and Gata6 in late blastocyst-stage (E5) rabbit embryos after microinjection of rhesus PSCs (confocal imaging; scale bars: 50 μm). (**D**) Percentage of GFP^+^/Nanog^+^, GFP^+^/Sox2^+^, and GFP^+^/Gata6^+^ cells in rabbit embryos 1–3 days after injection of rhesus PSCs (n = 232).

We subsequently characterized the surviving rhesus PSCs. The chimeric embryos were immunostained with GFP, OCT4, SOX2, NANOG, and GATA6 antibodies at E3 (1 DIV, 335 embryos with GFP^+^ cells, n = 650), E4 (2 DIV, 202 embryos with GFP^+^ cells, n = 664), and E5 (3 DIV, 217 embryos with GFP^+^ cells, n = 932). GFP^+^/OCT4^+^ and GFP^+^/GATA6^+^ co-immunostaining was frequently observed in chimeric embryos at 3 DIV. In contrast, GFP^+^/SOX2^+^ cells were rarely observed (**Fig. 3C**, **Table S1**). GFP^+^/SOX2^+^, GFP^+^/NANOG^+^, and GFP^+^/GATA6^+^ cells were counted in 754 positive embryos. A strong decrease in the rate of GFP^+^/SOX2^+^ and GFP^+^/NANOG^+^ cells and an increase in the rate of GFP^+^/GATA6^+^ cells were observed between 1 and 3 DIV with all reprogramming protocols (**Fig. 3D**). These results strongly suggest that the vast majority of rhesus PSCs abolished the expression of pluripotency markers and committed prematurely to differentiation after injection into rabbit embryos. A similar result was observed when rabbit embryos injected with TL2i rhesus PSCs were cultured for 3 days in gradual medium supplemented with 250 nM tamoxifen (**Supplementary data 9A**), indicating that the maintenance of high STAT3 activity is not sufficient to overcome the spontaneous differentiation that occurs during embryo colonization.

In previous experiments, we compared the colonization capabilities of mESCs and monkey PSCs using the same experimental paradigm, which included single-cell dissociation and embryo culture in 20% O_2_ concentration. Rabbit embryos are routinely cultured at atmospheric O_2_ concentration (Tapponnier et al., 2017). Thus, we compared the GFP^+^ cell colonization rates after culturing host embryos under normoxic (20% O2) and hypoxic (5% O2) conditions. Following injection of 10 rhesus TL2i cells, host embryos were cultured under conditions of 5% and 20% O_2_ concentration (number of embryos treated: 112 and 109, respectively). The rate of embryos containing GFP^+^ cells was lower at 1, 2, and 3 DIV after culture at 5% O_2_ concentration (**Supplemental data S9B**). We also evaluated whether microinjection of cell clumps into rabbit embryos would improve the colonization capabilities compared with single-cell suspensions. The experiment was performed using rhesus TL2i cells. At both 1 and 2 DIV, the rate of GFP^+^ embryos obtained after injection of small aggregates of 10 cells was lower than that obtained after injection of the same number of isolated cells (1 DIV: 37% *vs.* 53%, respectively; n = 76; 2 DIV: 19% *vs.* 43%, respectively; n = 76; **Supplemental data S9C**). In contrast, at 3 DIV, injection of aggregates of 10 cells yielded slightly better results than those recorded after injection of the same number of isolated cells (18.9% *vs.* 6.25% of GFP^+^ embryos, respectively; n = 98). However, we did not observe an increase in the number of GFP^+^ cells in the positive embryos. Finally, we evaluated the efficacy of colonization after microinjection of 10 cells into embryos at the morula stage (E2) compared with the blastocyst stage (E4) (**Supplemental data S9D**). At 3 DIV, we observed lower levels of GFP^+^ embryos for TL2i cells (39% *vs.* 6%) and 4i/L/b cells (29% *vs.* 0%) after microinjection into blastocyst-stage embryos. On the basis of these additional experiments, we concluded that injection of embryos at the blastocyst stage, injection of cell clumps, and reduction of O_2_ concentration during embryo culture had no measurable effects on the rate of embryo colonization by rhesus PSCs.

Previous studies have shown that inhibiting apoptosis either by the addition of the Rock inhibitor Y27632 (Kang et al., 2018), or by overexpression of the anti-apoptotic genes *BCL2* (Masaki et al., 2016) or BMI1 (Huang et al., 2018) enables primed PSCs to colonize host embryos. Therefore, in a final series of microinjections, we asked whether inhibition of apoptosis could improve the engraftment of naïve PSCs into the host embryo. For this purpose, NHSM-v and t2iLGöY PSCs were treated with the ROCK inhibitor Y27632 during dissociation and microinjection into the host embryos. The rates of embryos containing GFP^+^ cells were similar at all three time points analyzed (1, 2, and 3 DIV; **Supplemental data S9E**). Thus, addition of Rock inhibitor did not improve the rate of embryo colonization by rhesus naïve PSCs.

### Colonization of rabbit pre-implantation embryos by human induced PSCs

We subsequently investigated whether human naïve PSCs exhibited a similar behavior to that of rhesus naïve PSCs after injection into rabbit embryos. We tested the colonization capabilities of two human iPSC lines, cR-NCRM2 (Guo et al., 2017), and IR1.7 (Ng et al., 2012), after reprogramming with t2iLGöY (Guo et al., 2017) and TL2i protocols (Chen et al., 2015a), respectively. The cR-NCRM2-t2iLGöY cells express the naïve pluripotency markers TFAP2C, KLF4, KLF17, transcription factor CP2 like 1 (TFCP2L1), and sushi domain containing 2 (SUSD2) (Bredenkamp et al., 2019) (**Supplemental data S9F**). After injection into rabbit morulas, immuno-fluorescence was used for the detection of human nuclear antigen (HuN) to trace human cells in rabbit host embryos. On days 1, 2, and 3, 93 (n = 29), 65 (n = 31), and 33% (n = 130) of the embryos contained HuN^+^ cells in the ICM/epiblast, respectively (**Fig. 4C**, **Table S1**). At 3 DIV, double immunofluorescence analysis revealed that 45 (n = 12), 17 (n = 13), and 32% (n = 6) of the embryos contained HuN^+^/SOX2^+^, HuN^+^/NANOG^+^, and HuN^+^/GATA6^+^ cells, respectively (**Fig. 4A**, **D**). These rates are similar to those previously obtained using t2iLGöY rhesus PSCs (**Fig. 3B**), suggesting extremely similar colonization capabilities for rhesus and human t2iLGöY cells. Chimeric embryos also contained HuN^+^/SOX17^+^ (50%, n = 30) and HuN^+^/T-BRA^+^ cells (36%, n = 30), but none of the embryos examined contained HuN^+^/NESTIN^+^ cells (n = 30), suggesting preferential differentiation toward mesodermal and endodermal lineages (**Fig. 4A**, **D**). Similar results were obtained using naïve IR1.7-TL2i cells. These cells have been described previously (Chen et al., 2015a) (**Supplemental data S9F**). On days 1, 2, and 3, 85 (n = 39), 50 (n = 36), and 28% (n = 39) of embryos, respectively, contained HuN^+^ cells in the ICM/epiblast (**Fig. 4C**, **Table S1**). At 3 DIV, chimeric embryos contained HuN^+^/SOX2^+^ (10%, n = 76), HuN^+^/GATA6^+^ (36%, n = 70), and HuN^+^/SOX17^+^ (86%, n = 30) cells (**Fig. 4B**, **D**). None of the embryos examined contained HuN^+^/NANOG^+^ (n = 26), HuN^+^/T-BRA^+^ (n = 27), and HuN^+^/NESTIN^+^ (n = 30) cells. Considering all of the data obtained using both cR-NCRM2-t2iLGöY and IR1.7-TL2i cells, donor cells that colonized rabbit embryos expressed SOX2 (21%, n = 30), NANOG (8%, n = 5), GATA6 (32%, n = 33), SOX17 (58%, n = 35), and T-BRA (19%, n = 11) (**Fig. 4E**). These results illustrated both a low rate of colonization by SOX2^+^/NANOG^+^/OCT4^+^ human naïve PSCs at 3 DIV and premature differentiation of the donor cells to endodermal and mesodermal fates, thus confirming and extending the results obtained using rhesus naïve PSCs.

**Figure 4:**
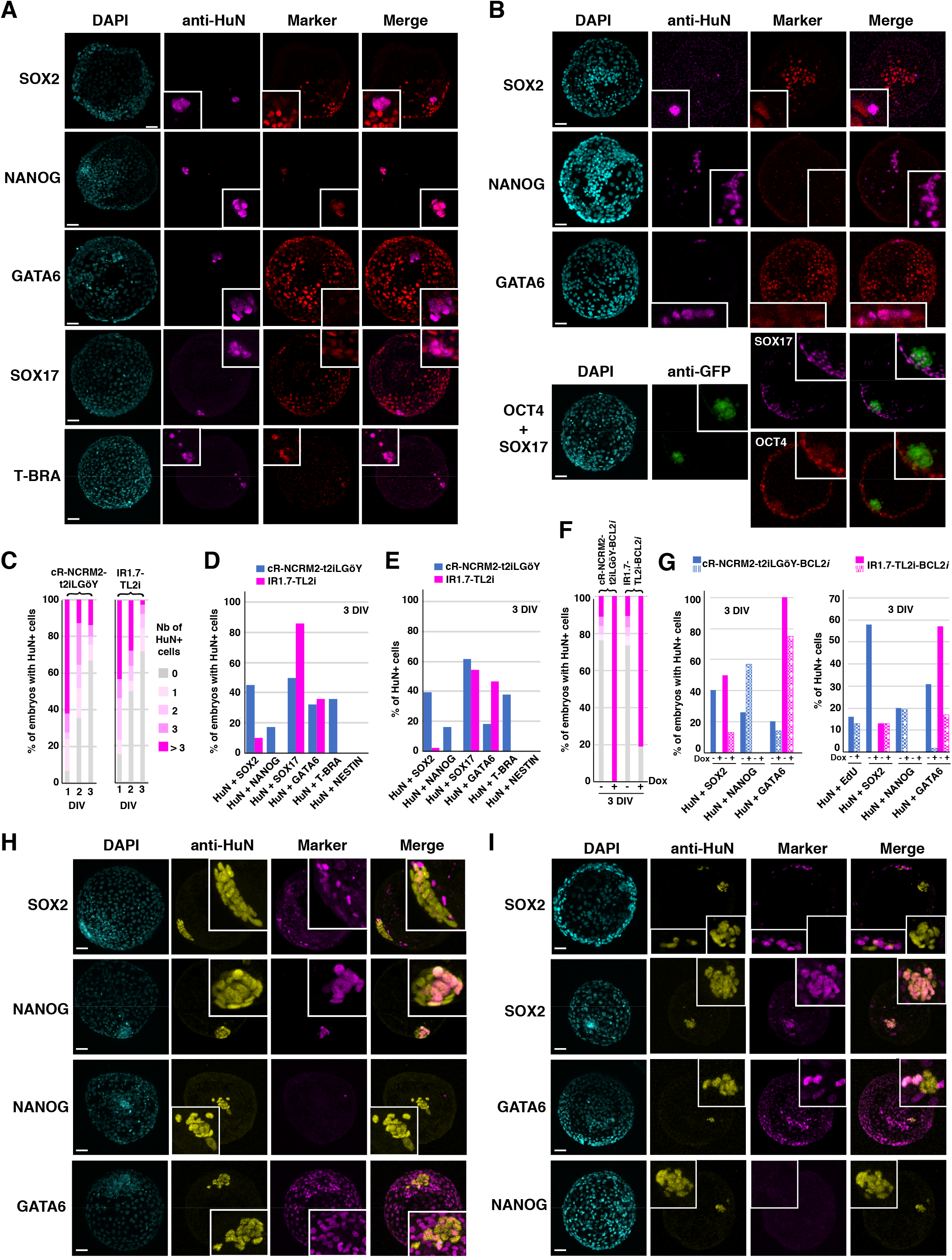
Colonization of rabbit embryos by human naïve PSCs. (**A**) Immunostaining of HuN, SOX2, NANOG, GATA6, SOX17, and T-BRA in late blastocyst-stage rabbit embryos (E5, 3 DIV) after microinjection of cR-NCRM2-t2iLGöY cells into morula-stage (E2) embryos (confocal imaging; scale bars: 50 μm). (**B**) Immunostaining of HuN, SOX2, NANOG, GATA6, OCT4 and SOX17 in late blastocyst-stage rabbit embryos (E5, 3 DIV) after microinjection of IR7.1-TL2i cells into morula-stage (E2) embryos (confocal imaging; scale bars: 50 μm). (**C**) Percentage of rabbit embryos with 0, 1, 2, 3, or >3 HuN^+^ cells at 1–3 days (1–3 DIV) after injection of cR-NCRM2-t2iLGöY (n = 191) and IR7.1-TL2i cells (n = 114). (**D**) Percentage of rabbit embryos with HuN^+^/SOX2^+^, HuN^+^/NANOG^+^, HuN^+^/GATA6^+^, HuN^+^/SOX17^+^, HuN^+^/T-BRA^+^, and HuN^+^/NESTIN^+^, cells at 3 DIV after injection of cR-NCRM2-t2iLGöY and IR7.1-TL2i cells. (**E**) Percentage of HuN^+^/SOX2^+^, HuN^+^/NANOG^+^, HuN^+^/OCT4^+^, HuN^+^/GATA6^+^, HuN^+^/SOX17^+^, HuN^+^/T-BRA^+^, and HuN^+^/NESTIN^+^ cells at 3 DIV after injection of cR-NCRM2-t2iLGöY and IR7.1-TL2i cells. (**F**) Percentage of rabbit embryos with 0, 1, 2, 3, or >3 HuN^+^ cells at 3 days (3 DIV) after injection of cR-NCRM2-t2iLGöY-BCL2*i* (+/- Dox), and IR7.1-TL2i-BCL2*i* cells (+/- Dox). (**G**) Left panel: percentage of rabbit embryos with HuN^+^/SOX2^+^, HuN^+^/NANOG^+^, and HuN^+^/GATA6^+^ cells at 3 DIV after injection of cR-NCRM2-t2iLGöY-BCL2*i* and IR7.1-TL2i-BCL2*i* cells (+/- Dox); right panel: percentage of HuN^+^/EdU^+^, HuN^+^/SOX2^+^, HuN^+^/NANOG^+^, and HuN^+^/GATA6^+^ cells at 3 DIV after injection of cR-NCRM2-t2iLGöY-BCL2*i* and IR7.1-TL2i-BCL*i* cells (+/- Dox). (**H**) Immunostaining of HuN, SOX2, NANOG, and GATA6 in late blastocyst-stage rabbit embryos (E5, 3 DIV) after microinjection of cR-NCRM2-t2iLGöY-BCL2*i* cells into morula-stage (E2) embryos (confocal imaging; scale bars: 50 μm). (**I**) Immunostaining of HuN, SOX2, NANOG, and GATA6 in late blastocyst-stage rabbit embryos (E5, 3 DIV) after microinjection of IR7.1-TL2i-BCL2*i* cells into morula-stage (E2) embryos (confocal imaging; scale bars: 50 μm).

We then asked whether the inhibition of apoptosis via overexpression of the anti-apoptotic gene *BCL2* (Masaki et al., 2016) could improve the engraftment of human naïve iPSCs and prevent their premature differentiation. Both cR-NCRM2-t2iLGöY and IR1.7-TL2i cells were transfected with doxycycline (Dox)-inducible human BCL2. Stable transfectants expressing BCL2 in response to Dox were selected in neomycin (hereafter called cR-NCRM2-t2iLGöY-BCL2*i* and IR1.7-TL2i-BCL2*i* cells; **Supplemental data S9G**) and subsequently injected into rabbit morulas prior to culture for 3 DIV in Dox- or vehicle (control)-supplemented gradual medium. In Dox-supplemented medium, 100 (n = 82) and 81% (n = 76) of the embryos injected with cR-NCRM2-t2iLGöY-BCL2*i* and IR1.7-TL2i-BCL2*i* cells, respectively, exhibited the presence of HuN^+^ cells in the ICM/epiblast stage, whereas these rates were only 30 (n = 66, p < 0.001, χ^2^ test) and 34% (n = 77, p < 0.001, χ^2^ test), respectively, for control cells (**Fig. 4F, Table S1**). Moreover, at 3 DIV, the Dox-induced embryos contained more HuN^+^ cells than the control embryos (−Dox: 3.3 ± 1.55 cells/embryo; +Dox: 18.7 ± 5.5 cells/embryo; p < 0.001, Student’s *t*-test; **Fig. 4H-I**). These results strongly suggest that BCL2 overexpression prevents donor cell death after embryo injection. It should be noted that the percentage of EdU^+^ cells was not altered [−Dox: 16% (n = 20), +Dox: 13% (n = 20); p < 0.69, χ^2^ test; **Figs. 4G and S9H**], indicating that BCL2 overexpression had no measurable effect on the mitotic index of the donor cells. The identity of the surviving HuN^+^ cells was subsequently analyzed via double immunofluorescence at 3 DIV. In the +Dox condition, most embryos contained HuN^+^/GATA6^+^ cells, suggesting that BCL2 overexpression does not alter the natural propensity of donor cells to differentiate after injection into host embryos (**Fig. 4G– I**). To eliminate the confounding effect of the higher HuN^+^ cell count observed in Dox-treated embryos, we counted the number of HuN^+^ cells expressing SOX2, NANOG, and GATA6 in both induced (+Dox; n = 144) and uninduced embryos (−Dox; n = 143). The proportions of HuN^+^/GATA6^+^ cells were strongly diminished after BCL2 induction [2% (n = 27) *vs.* 31% (n = 20) for cR-NCRM2-t2iLGöY-BCL2*i* cells, p < 0.001, χ^2^ test; 17% (n = 25) *vs.* 57% (n = 25) for IR1.7-TL2i-BCL2*i* cells, p < 0.001, χ^2^ test]. However, the sharp decrease in the proportion of HuN^+^/GATA6^+^ cells was not associated with mirror increases in the proportions of HuN^+^/NANOG^+^ [19.5% (n = 27) *vs.* 20% (n = 26) for cR-NCRM2-t2iLGöY-BCL2*i* cells; 0% (n = 52) for IR1.7-TL2i-BCL2*i* cells in both +Dox and −Dox conditions] and HuN^+^/SOX2^+^ cells [13% (n = 25) in both +Dox and −Dox conditions]. On the contrary, it was strongly diminished in cR-NCRM2-t2iLGöY-BCL2*i* cells [0% (n = 28) *vs.* 58% (n = 20), p < 0.001, χ^2^ test]. Thus, BCL2 overexpression significantly increased the efficiency of embryo colonization by human naïve PSCs. However, in most of the examined embryos, the vast majority of the donor cells failed to maintain the expression of pluripotency markers, and many of them remained unspecified or expressed GATA6.

### Colonization of cynomolgus monkey pre-implantation embryos by rhesus PSCs and human iPSCs

We questioned whether the low rate of proliferation and ICM/epiblast colonization of human TL2i and t2iLGöY cells after injection into rabbit morulas were due to the evolutionary distance between the two species. To answer this question, we injected human cR-NCRM2-t2iLGöY and IR1.7-TL2i cells, and rhesus TL2i cells into cynomolgus monkey embryos. Ten cells of each were injected into embryos at the morula stage (E4). The embryos were subsequently cultured for 3 DIV in gradual media and developed into mid/late (E7) blastocysts before being subjected to triple immunostaining for GFP, OCT4, and NANOG. Of the cynomolgus embryos, 13% (n = 15) had incorporated human cR-NCRM2-t2iLGöY cells, for a total of 5 GFP^+^ cells (2 and 3 GFP^+^ cells/embryo), of which only 1 was both OCT4^+^ and NANOG^+^ (**Fig. 5A**, **Table S1**); 29% (n = 7) of the cynomolgus embryos had incorporated human IR1.7-TL2i cells, for a total of 14 GFP^+^ cells (10 and 4 GFP^+^ cells/embryo), of which 2 were either OCT4^+^ or NANOG^+^ (**Fig. 5B**, **Table S1**); and 29% (n = 7) of cynomolgus embryos had incorporated rhesus TL2i cells, for a total of 3 GFP^+^ cells (1 and 2 GFP^+^ cells/embryo), none of which were either Oct4^+^ or NANOG^+^ (**Fig. 5C**, **Table S1**). These results indicate that human iPSCs (IR1.7-TL2i and NCRM2-t2iLGöY) and rhesus monkey PSCs [LyonES-tGFP-(S3)-TL2i] show similarly behaviors after injection into closely and distantly related embryos (cynomolgus and rabbit embryos, respectively).

**Figure 5:**
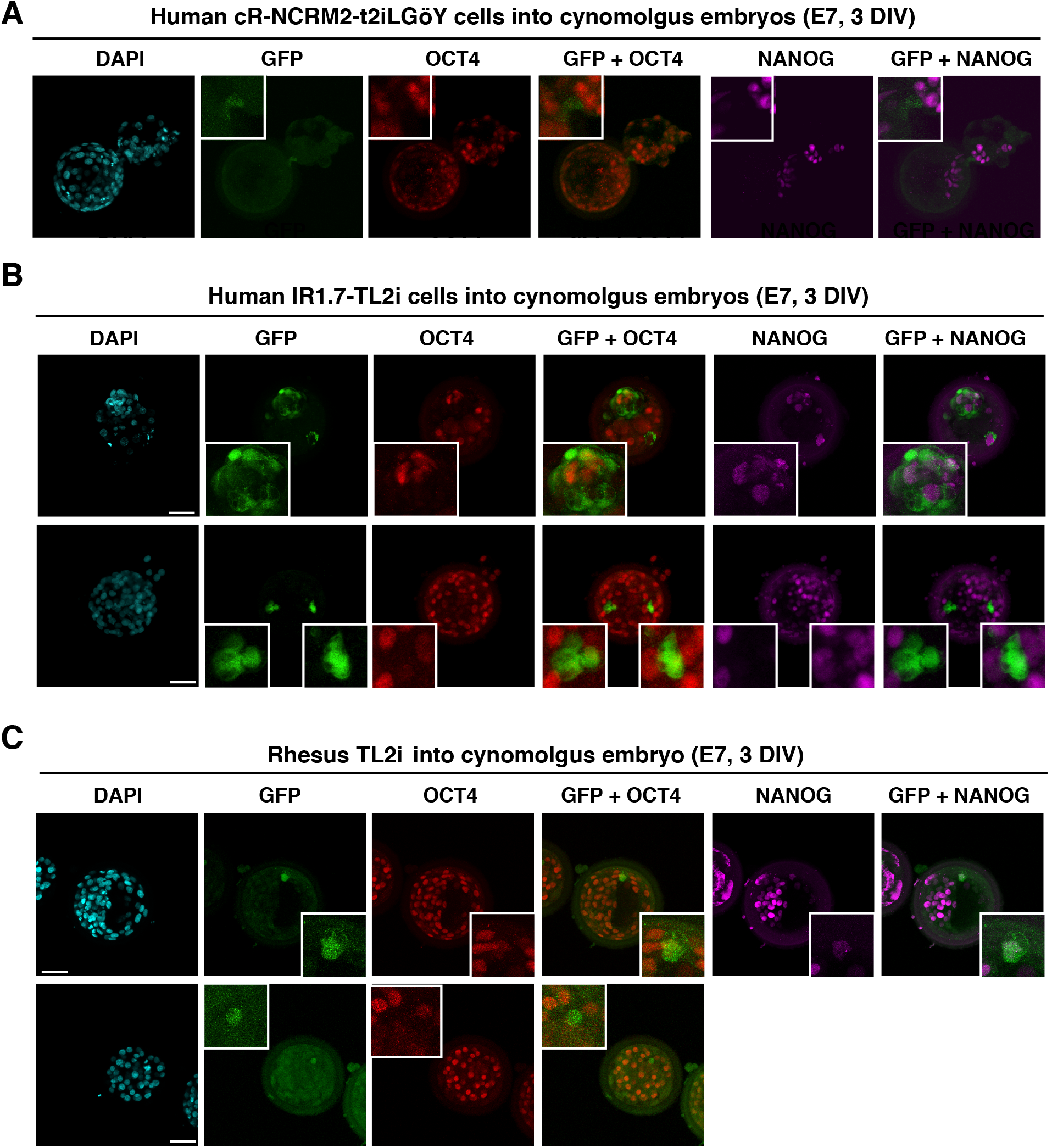
Colonization of cynomolgus monkey embryos by human iPSCs and rhesus PSCs. (**A**) Immunolabeling of GFP, OCT4, and NANOG in late blastocyst-stage cynomolgus embryos (E7) after microinjection of human NCRM2-t2iLGöY cells. (**B**) Immunolabeling of GFP, OCT4, and NANOG in late blastocyst-stage cynomolgus embryos (E7) after microinjection of human IR7.1-TL2i cells. (C) Immunolabeling of GFP, OCT4, and NANOG in late blastocyst-stage cynomolgus embryos (E7) after microinjection of rhesus TL2i cells. (**A-C**) Confocal imaging; scale bars: 50 μm).

### Contrasting cell-cycle parameters of mESCs, rhesus and human naïve PSCs

To understand why rhesus and human PSCs stop multiplying when injected into host embryos, we examined both the DNA replication and cell-cycle distribution. We started by examining the incorporation of EdU as a measure of the fraction of cells that replicate in the cell population (% EdU^+^). Measurements were performed in adherent cells (“Adh”), after single-cell dissociation with trypsin (“0h”), and after re-incubation for 1 hour (“1h”) and 2 hours (“2h”, human cells only) at 37°C in order to mimic the conditions encountered by the cells during injection into host embryos. mESCs incorporate EdU very rapidly (**Fig. 6A**), consistent with previous results describing the high rate of 5-bromo-2-deoxyuridine (BrdU) incorporation in mESCs (Savatier et al., 1996). Single-cell dissociation did not alter the percentage of EdU^+^ mESCs (77% and 78% of EdU^+^ cells in “Adh” and “0h” cells, respectively). It slightly decreased in “1h” cells (48% of EdU^+^ cells); however, the rate of EdU incorporation remained unchanged. These observations suggest that DNA replication is not altered by the experimental setting in the vast majority of mESCs. Similar conclusions were made with mESCs cultured in 2i/LIF medium. However, it should be noted that the rate of EdU incorporation was lower under 2i/LIF culture conditions compared with that under serum/LIF conditions. This finding is consistent with those of a previous report showing that mESCs cultured in 2i/LIF exhibit a slower cell cycle compared with their serum/LIF counterparts (Ter Huurne et al., 2017).

**Figure 6:**
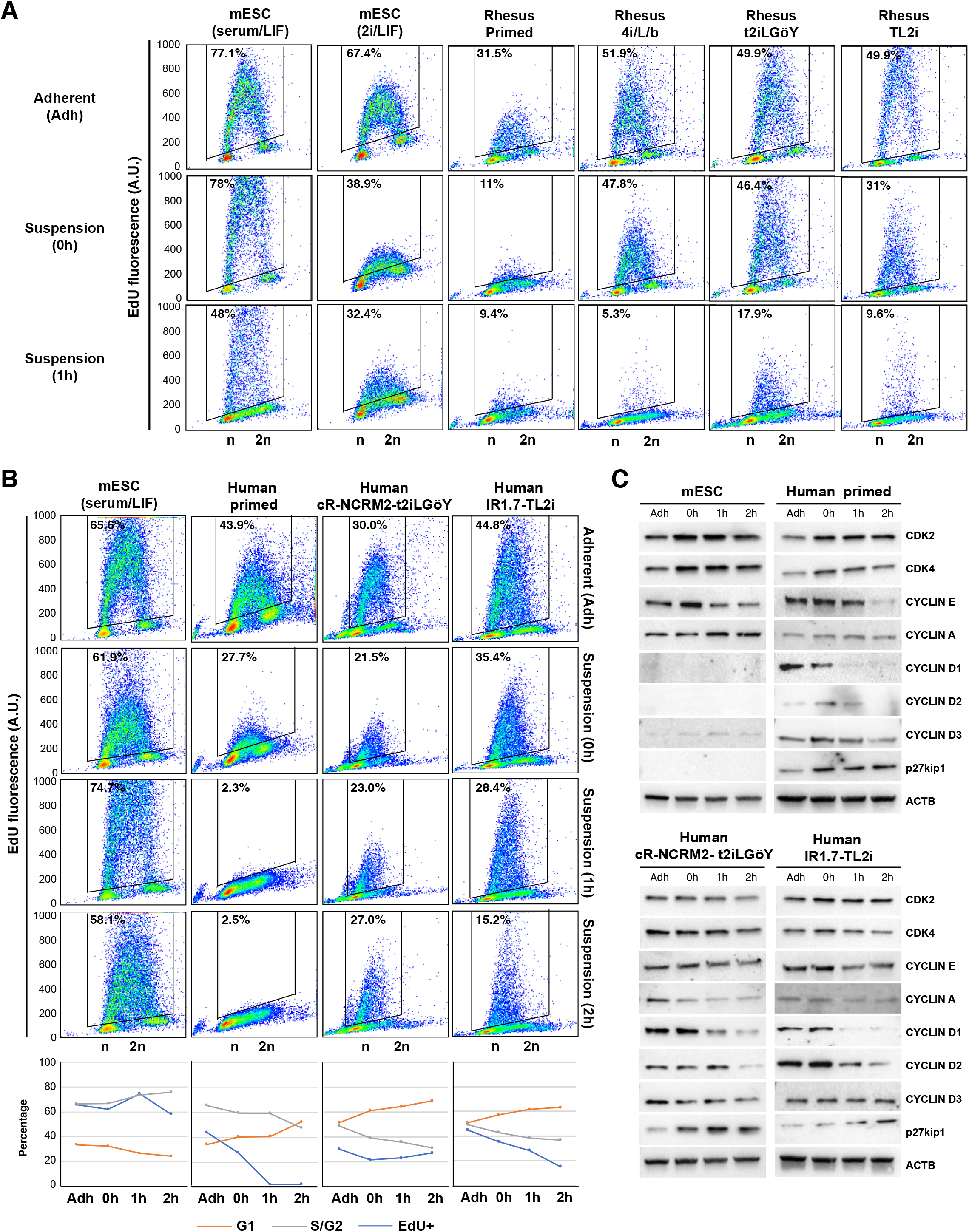
Cell-cycle parameters of mouse ESCs, rhesus PSCs, and human iPSCs, before and after reprogramming to naïve states. (**A**) Flow cytometry analysis of EdU incorporation and propidium iodide (PI) staining of mESCs (serum/LIF and 2i/LIF) and rhesus PSCs (primed, 4i/L/b, TL2i, and t2iLGöY) in “Adh”, “0h”, and “1h” condition. Numbers indicate percentages of EdU^+^ cells. (**B**) Flow cytometry analysis of EdU incorporation and PI staining of mESCs (serum/LIF) and human iPSCs (primed, cR-NCRM2-t2iLGöY and IR7.1-TL2i) in “Adh”, “0h”, “1h”, and “2h” conditions. Numbers indicate percentage of EdU^+^ cells. Histograms show the cell cycle distribution (% EdU^+^, G1, and S/G2 cells). (**C**) Western blot analysis of cell-cycle regulators in mESCs (serum/LIF) and human iPSCs (primed, cR-NCRM2-t2iLGöY and IR7.1-TL2i) in “Adh”, “0h”, “1h”, and “2h” condition (results of 3 experiments).

The pattern of EdU incorporation in primed LyonES-tGFP-(S3) cells was markedly different. The percentage of EdU^+^ cells and the incorporation rate were significantly reduced in “Adh” cells compared to those measured in mESCs (**Fig. 6A**). They were further reduced in “0h” and “1h” cells. In “1h” cells, DNA replication was almost completely abolished. This reveals the inherent failure of primed LyonES-tGFP-(S3) cells to sustain cell cycle progression following single-cell dissociation. Interestingly, reprogramming primed LyonES-tGFP-(S3) cells with 4i/L/b, TL2i and t2iLGöY protocols significantly increased the percentage of EdU^+^ cells, consistent with a previous report indicating that resetting naïve-like features accelerated the cell cycle of human PSCs (Chen et al., 2015a). However, unlike mESCs, rhesus naïve 4i/L/b, TL2i and t2iLGöY cells slowed DNA replication after dissociation, evidenced by a significant reduction of EdU incorporation in both ”0h” and “1h” cells. In “1h” cells, only 6%, 10%, and 18% of the 4i/L/b, TL2i and t2iLGöY cells, respectively, incorporated EdU (compared to 48% of mESCs in the same experimental condition). Furthermore, the incorporation rate was considerably reduced as compared to “Adh” 4i/L/b, TL2i and t2iLGöY cells and “1h” control mESCs. Overall, these results indicate that rhesus naïve PSCs, unlike mESCs, fail to maintain active DNA replication in this experimental setting. In addition, the cell cycle distribution of the “1h” samples showed a slight increase in the proportion of cells in the G1 phase at the expense of cells in the G2 phase in rhesus primed 4i/L/b, TL2i and t2iLGöY cells, suggesting that some cells have already undergone growth arrest in G1 phase. This alteration was not observed in mESCs (**Supplemental data S10A**). Taken together, these results indicate that the vast majority of naïve rhesus cells are markedly slowed down in their mitotic cycle at the time they are injected into host embryos, or shortly after.

Similar results were obtained using human IR1.7-TL2i and cR-NCRM2-t2iLGöY cells in the “Adh,” “0h,” and “1h” conditions. A “2h” condition (i.e., culture in suspension for 2 h) was added to the experimental scheme. The percentage of EdU^+^ cells was reduced under the “0h,” “1h,” and “2h” cells compared with the findings in “Adh” cells (**Fig. 6B**). In “2h” cells, only 15.2% of the IR1.7-TL2i cells incorporated EdU (compared with 43.8% in “Adh” IR1.7-TL2i cells, 2.5% in primed cells, and 58.1% in “2h” mESC). The impact of suspension culture was less pronounced in human cR-NCRM2-t2iLGöY cells. However, the rate of EdU incorporation was significantly lower in cR-NCRM2-t2iLGöY cells than in IR1.7-TL2i cells. Interestingly, human cR-NCRM2-t2iLGöY cells exhibited a unique pattern of EdU incorporation, in which the percentage of EdU^+^ cells was higher at “2h” (27%) than at “0h” (21.5%). This observation suggests that some cells in the human cR-NCRM2-t2iLGöY population may have the capacity to recover from the deleterious effects of single-cell dissociation and gradually re-engage in the mitotic cycle. This phenomenon was not observed in any other cell lines. We also analyzed the cell cycle distribution (**Fig. 6B** and **Supplemental data S10B**). In all human cells (primed, IR1.7-TL2i and cR-NCRM2-t2iLGöY), the cell cycle distribution of the “2h” samples exhibited an increased proportion of cells in G1 phase at the expense of cells in S and G2 phases, suggesting that some cells have already undergone growth arrest in G1 phase. This alteration was not observed in mESCs, further indicating that only mESCs retain the capacity to actively divide in non-adherent culture conditions.

To understand why the mitotic cycle is slowed in human naïve PSCs after single-cell dissociation, we studied the expression of cell cycle regulators, including CDK2, CDK4, cyclins D1, D2, D3, E, and A, and the CDK inhibitor p27kip1 (**Fig. 6C** and **Supplemental data S10C**). No differences of CDK2 and CDK4 levels were observed among “Adh,” “0h,” “1h,” and “2h” samples in all cell types analyzed. In mESCs, the levels of cyclins E and A were increased. Cyclin D1, cyclin D2, and p27kip1 were not expressed as previously reported (Savatier et al., 1996) (Stead et al., 2002). In human primed PSCs, the expression of cyclins D1, D2, and D3 was decreased and that of p27kip1 was increased after dissociation and suspension culture. In cR-NCRM2-t2iLGöY cells, the expression of cyclins D1, D2, D3, and A was decreased, and that of p27kip1 was increased. In IR1.7-TL2i cells, the expression of cyclins D2 and A was decreased, and that of p27kip1 was increased. Therefore, single-cell dissociation followed by suspension culture triggers opposite responses in mESCs and human naïve PSCs. in mESCs, it activates the expression of positive regulators of the mitotic cycle, whereas in human naïve PSCs, it reduces their levels, and concomitantly increases that of a negative regulator. These results are consistent with the results of EdU incorporation and cytometry analyses, revealing cell cycle delay, and possibly growth arrest in G1 phase.

### Rhesus PSCs stall in the G1 phase after transfer into rabbit embryos

We observed that mESCs continue to actively replicate their DNA after injection into rabbit embryos (**Fig. 1C**). Therefore, we investigated whether naïve rhesus cells that survive injection and successfully colonize the ICM/epiblast retain a high proliferative capacity. For this purpose, 10 rhesus TL2i cells were injected into rabbit embryos at the morula stage. The embryos were subsequently cultured for 1 and 2 DIV prior to pulse-labeling with EdU. At 1 DIV, seven embryos had GFP^+^ cells in the ICM/epiblast (37%, n = 19) for a total of 29 GFP^+^ cells. Triple immunostaining for GFP, SOX2, and EdU showed that 24 (82%) of these 29 GFP^+^ cells expressed SOX2, of which only four have incorporated EdU (**Supplemental data S11**). At 2 DIV, nine embryos had GFP^+^ cells in the ICM/epiblast (60%, n = 15) for a total of 61 GFP^+^ cells. Of these 61 GFP^+^ cells, 44 (72%) expressed SOX2, and 1 incorporated EdU (2%). These results reveal that the rhesus naïve cells that survived and colonized the rabbit embryos retarded their DNA replication and potentially underwent growth arrest.

To obtain further insight into the cell cycle of the mESCs and rhesus 4i/L/b, TL2i, and t2ILGöY naïve PSCs after injection into rabbit embryos, we generated cell lines expressing the FUCCI(CA) reporter system (Sakaue-Sawano et al., 2017). The mESCs expressing *PB-Puro^R^-CAG-mVenus:hGeminin-IRES-mCherry:hCdt1*, hereinafter referred to as mESC-FUCCI(CA), exhibited a cell cycle distribution typical of mESCs (18%, 69%, and 11% of cells in G1, S, and G2 phases, respectively; **Supplemental data S12A, B**). On the other hand, rhesus PSC-FUCCI(CA) cells exhibited an extended G1 phase relative to other cell cycle phases (43%, 36%, and 13% of the cells in G1, S, and G2 phases, respectively) (**Supplemental data S12C, D**). After injection into rabbit embryos, mESC-FUCCI(CA) cells proliferated actively during the 3 DIV, as shown by the expansion of the mVenus^+^ cell population identified through immunostaining with an anti-mVenus antibody (green) in 100% of the embryos (**Figs. 7A, B**). Even after 3 DIV, mCherry^+^ cells (red), identified using an anti-mCherry antibody, and mVenus^+^/mCherry^+^ cells (yellow) were proportionally less numerous, indicating that only rare mESCs were in G1 and G2 phases, respectively. In addition, at all time-points analyzed, there were no embryos containing only mCherry^+^ (red) cells (n = 27) (**Fig. 7B**). These results indicate that mESCs continue to proliferate actively for at least three days after injection into host embryos, in line with the EdU incorporation data.

**Figure 7:**
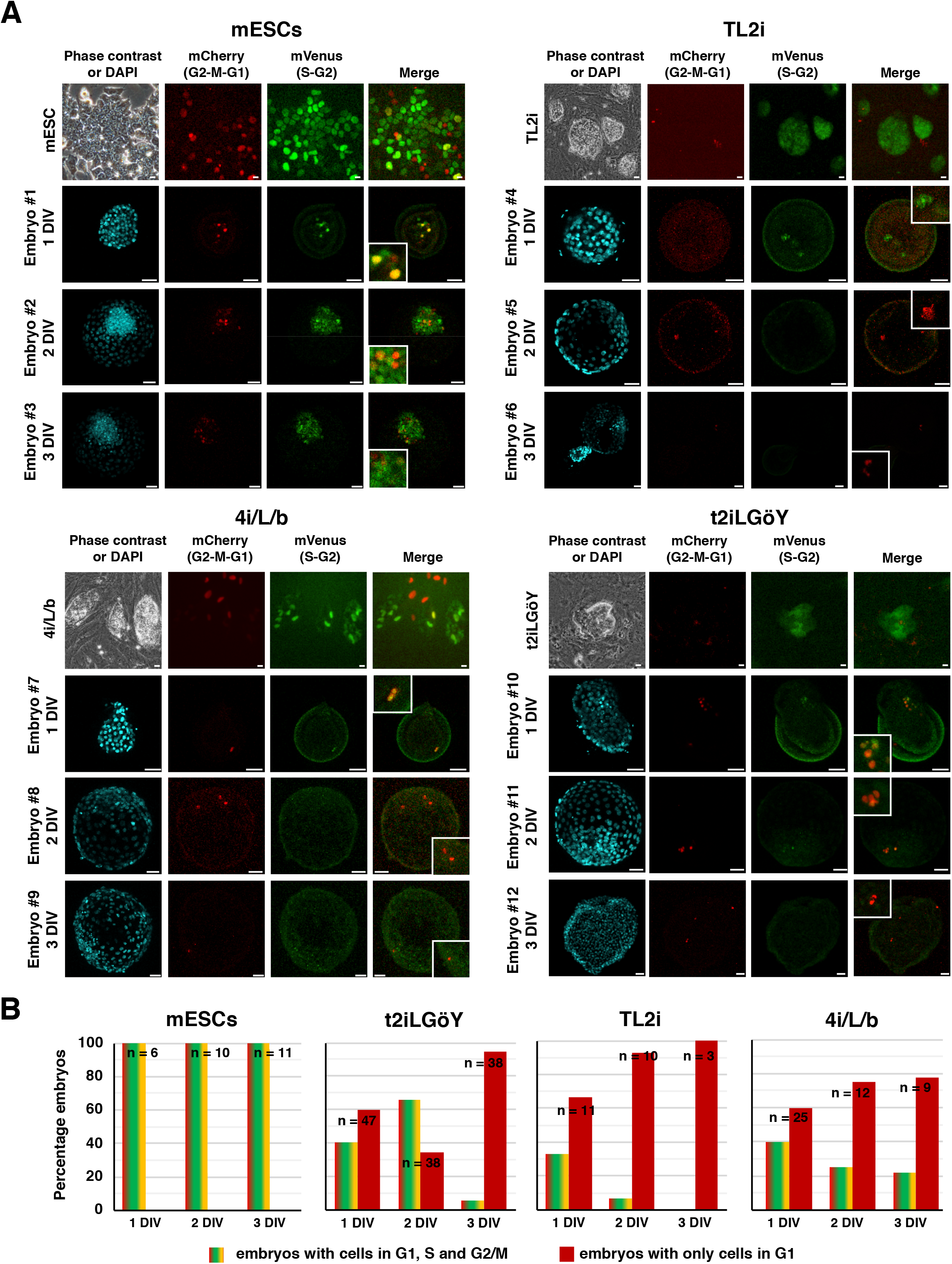
Cell cycle distribution of mouse and rhesus PSCs after injection into rabbit embryos. (**A**) Phase contrast and epifluorescence imaging of mouse ESCs and rhesus TL2i, 4i/L/b, and t2iLGöY PSCs expressing the FUCCI(CA) cell cycle reporter system. Immunostaining of mVenus and mCherry in rabbit embryos 1–3 days (1–3 DIV) after microinjection of FUCCI(CA) mouse and rhesus PSCs into morula-stage (E2) embryos (confocal imaging, scale bars: 50 μm). **(B)** Histogram of the percentage of rabbit embryos having cells in G1, S, and G2 phase of the cell cycle at 1, 2, and 3 DIV (n = 220).

On the other hand, the cell cycle distribution of rhesus PSC-FUCCI(CA) cells after injection into rabbit embryos was clearly different, regardless of the method used (4i/L/b, TL2i, and t2iLGöY) for reprogramming to the naïve state. At 1 DIV, the majority of embryos contained only mCherry^+^ (red) cells (t2iLGöY, 60%, n = 47; TL2i, 68%, n = 11; 4i/L/b, 60%, n = 25). The remaining embryos contained both mVenus^+^ (green) and mVenus^+^/mCherry^+^ (yellow) cells in addition to mCherry^+^ (red) cells. In embryos injected with TL2i and 4i/L/b cells, the rate of embryos containing only mCherry^+^ (red) cells increased over time in culture, reaching 78% (n = 9) and 100% (n = 3) with 4i/L/b and TL2i cells, respectively, at 3 DIV (**Fig. 7B**). In embryos injected with t2iLGöY cells, the proportion of embryos containing only mCherry^+^ (red) cells first decreased at 2 DIV (35%, n = 36) and subsequently increased at 3 DIV (95%, n = 35) **(Fig. 7B)**. These results indicate that most naïve rhesus PSC-FUCCI(CA) cells accumulate in the G1 phase of the cell cycle after injection into rabbit embryos.

## Discussion

In our study, we aimed to understand why human naïve PSCs failed to colonize mouse and pig embryos (Masaki et al., 2015; Theunissen et al., 2016; Wu et al., 2017), despite exhibiting characteristic features of naïve pluripotency as primarily defined in rodents. Our main conclusion is that human and NHP naïve PSCs are inherently unfit to remain mitotically active during embryo colonization and therefore differentiate prematurely.

The first question we addressed was whether or not host embryos are permissive to colonization by naïve PSCs. To answer this question, we injected mouse ESCs into rabbit and cynomolgus embryos and showed that they continue to express pluripotency markers, they remain mitotically active for at least 3 days *in vitro*, and they contributed to the expansion of the host epiblast. Some of them contributed to the formation of the neural tube in rabbit post-implantation embryos. We conclude that the ICM and epiblast of rabbit and cynomolgus embryos are permissive to colonization by bona fide naïve PSCs. These results may seem to contradict those of Wu *et al.* who injected mouse ESCs into pig embryos and observed no chimerism at d21-d28 of gestation (Wu et al., 2017). Differentiated cells derived from mouse ESCs are likely to disappear from the chimera through cell competition in later stages of development, however, this should not obscure the fact that the mouse ESCs are initially capable of colonizing pre-implantation embryos very efficiently.

In sharp contrast with mESCs, none of the naïve rhesus and human PSCs tested in the same experimental paradigm is capable of massively colonizing the ICM and epiblast of the host, whether rabbit or cynomolgus monkey embryos. Some cells retaining the expression of pluripotency markers were observed at 3 DIV in rabbit embryos after injection of LCDM (EPS) (9% of total embryos), 4i/L/b (2% of total embryos), TL2i (5% of total embryos), TL-CDK8/19i (2% of total embryos), and t2iLGöY PSCs (3% of total embryos) (Table 1). In these 234 chimeric embryos, the total number of GFP^+^ cells expressing pluripotency markers after 3 DIV was 3.26 +/−1.27, when 10 cells were injected at the morula stage. The other cells differentiated prematurely. In contrast, up to 40 GFP^+^/NANOG^+^ and 36 GFP^+^/SOX2^+^ cells/embryo were observed following injection of 10 mESCs into rabbit morulas after the same period of culture. Similarly, up to 23 GFP^+^/SOX2^+^ cells/embryo were observed after injection of mESC-2i/LIF. These results illustrate the striking difference observed between mouse ESCs and primate PSCs in the stability of pluripotency gene expression after injection into host embryos. As a result, none of the protocols tested confer an embryo colonization competence to primate naïve PSCs similar to that of mESCs. It should be noted however that the different protocols tested for reprogramming human and NHP PSCs did not yield identical results. We were unable to generate undeniable chimeric blastocysts with the E-NHSM and NHSM-v protocols. This result is in contradiction with a previous publication reporting efficient colonization of cynomolgus monkey embryos by PSCs reprogrammed using the NHSM-v protocol (Chen et al., 2015b). This difference can be explained by the immunodetection of GFP^+^ cells in the developing embryos, which eliminates the confounding effect of autofluorescence due to dying and necrotic cells. We did not observe significant differences in the rate of colonization by GFP^+^/NANOG^+^, GFP^+^/Oct4^+^, and SOX2^+^/GFP^+^ cells between the other five reprogramming protocols. We do not rule out the possibility that some of these protocols may be more effective than others in generating cells capable of colonization. However, the number of unquestionable chimeric blastocysts obtained with each of these protocols was insufficient to conclude about significant differences.

After injection into rabbit and macaque monkey morulas, most rhesus and human naïve PSCs undergo cell death. We showed that the few cells that survive after 3 DIV stall in the G1 phase and commit to differentiation evidenced by the loss of pluripotency markers (OCT4, SOX2 and NANOG) and the activation of differentiation markers (GATA6, SOX17, and T-BRA). Cell death can be prevented by overexpressing BCL2, leading to an apparent increase in the rate of colonization by donor cells. However, the proportion of pluripotent cells remains globally unchanged, indicating that the prevention of cell death does not prevent the premature differentiation observed with BCL2 transgene expression-negative cells. The behavior of primate naïve PSCs is in sharp contrast with that of mESCs, which continue to actively proliferate after injection into host embryos. These results reveal the inability of human and rhesus monkey naïve PSCs to remain mitotically active in an unfavorable environment and question their very nature. Despite requiring complex culture media, human and NHP naïve PSCs self-renew in a more precarious equilibrium than mESCs. This suggests a much higher sensitivity to alterations of their local environment, possibly explaining the poor performance of human and rhesus monkey naïve PSCs in colonizing the epiblast of host embryos. As a matter of fact, disruption of the human and rhesus PSC cell cycle is already evidenced immediately after the colony dissociation (i.e. prior to embryo injection) where most naïve PSCs cells undergo a drastic slowdown in DNA replication. Mitotic arrest is associated with down-regulation of cyclin expression and an up-regulation of p27kip1 expression. We speculate that only few cells retain the capacity to actively divide and re-enter the next division cycle when introduced into the embryo environment.

In the prospect of generating interspecies chimeras, an equally important issue is the influence of the host species on the rate of colonization. Rhesus and human PSCs (TL2i and t2iLGöY reprogramming protocols) did not show a marked difference in survival rate and stability of pluripotency gene expression after injection into rabbit and cynomolgus embryos. Similarly, mouse ESCs were able to colonize both rabbit and cynomolgus embryos with high efficiency. This suggests that the rabbit embryo is a valuable model system for exploring further improvements.

## Supporting information

Supplemental Experimental Procedures

## Experimental Procedures

A detailed description of the procedures is provided in Supplemental Experimental Procedures.

## Data and code availability

The RNA-Seq datasets generated during this study are available under GEO accession number: GSE146178.

## Acknowledgments

We are grateful to all members of the animal facility team for their work and dedication, and to Charlotte Bréhier for technical help during the revision of the manuscript. We thank Dr. Austin Smith for sharing the cR-NCRM2 cell line and RIKEN BRC DNABank for providing the mCherry-hCdt1(1/100)Cy(-)/pcDNA3 and mVenus-hGeminin(1/110)/pcDNA3 plasmids. This work was supported by the *Fondation pour la Recherche Medicale* (DEQ20170336757 and ARF20140129246), The *Fondation pour la recherche contre le cancer* (RAC18005CCA), the *Infrastructure Nationale en Biologie et Santé* INGESTEM (ANR-11-INBS-0009), the IHU-B CESAME (ANR-10-IBHU-003), the LabEx REVIVE (ANR-10-LABX-73), the LabEx “DEVweCAN” (ANR-10-LABX-0061), the LabEx “CORTEX” (ANR-11-LABX-0042), and the University of Lyon within the program “*Investissements d'Avenir*”(ANR-11-IDEX-0007). Work in the laboratory of M.S. was funded by the IRB and by grants from the Spanish Ministry of Economy co-funded by the European Regional Development Fund (ERDF) (SAF2017-82613-R), the European Research Council (ERC-2014-AdG/669622) and “La Caixa” Foundation.

## Authors’ contribution

I.A., C.R., A.M., E.M., V.C., A.B-M., N.D., F.W., P-Y.B., and M.A. performed cell and embryo cultures and cell characterizations.

M.D. and T.J. performed animal surgery.

G.M., C.M., and O.R. performed bioinformatic analyses.

C.L., M.S., and C.D. shared unpublished expertise.

I.A. and P.S. analyzed the data and wrote the manuscript.

## Supplemental data movie legends

**Movie S2**: Two-photon microscopy analysis of mESC-GFP cells after microinjection into a rabbit embryo (E2) and follow-up for 48–72 h.

**Movie S5:** Two-photon microscopy analysis of rhesus 4i/L/b PSCs after microinjection into a rabbit embryo (E2) and follow-up for 48–72 h.

**Movie S6:** Two-photon microscopy analysis of rhesus t2iLGöY PSCs after microinjection into a rabbit embryo (E2) and follow-up for 48–72 h.

**Movie S7:** Two-photon microscopy analysis of Rhesus TL2i PSCs after microinjection into a rabbit embryo (E2) and follow-up for 48–72 h.

**Movie S9:** Two-photon microscopy analysis of Rhesus E-NHSM PSCs after microinjection into a rabbit embryo (E2) and follow-up for 48–72 h.

