## Supplemental Experimental Procedures for "Apoptosis, G1 phase stall and premature differentiation account for low chimeric competence of Human and rhesus monkey naïve pluripotent stem cells"

**Inventory of Supplemental Information**

Figure S1

Figure S3

Figure S4

Figure S9

Figure S10

Figure S11

Figure S12

Table S1

Table S3

Table S4

Supplemental Experimental Procedures

Supplemental References

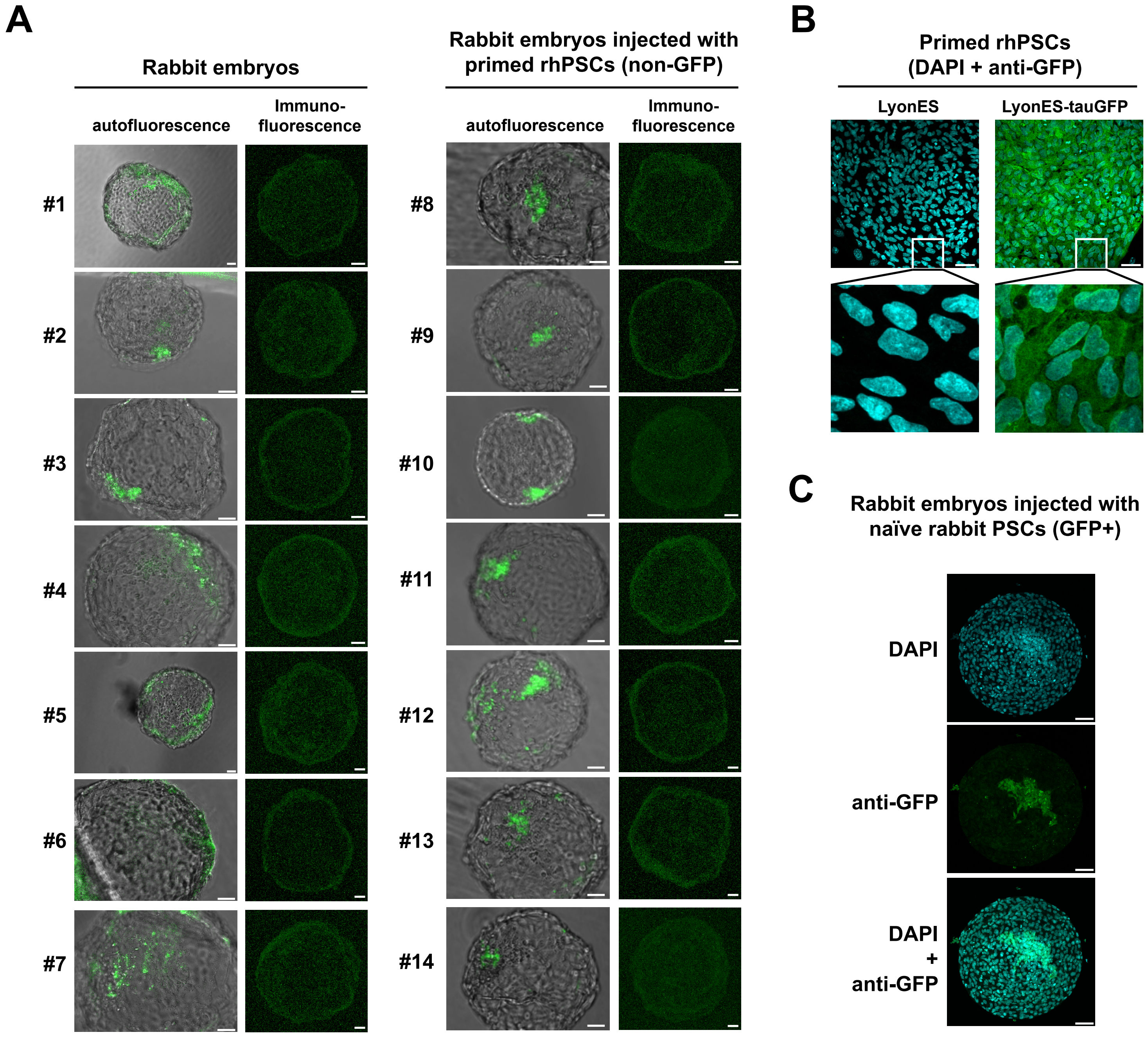

**Figure S1: Evaluation of autofluorescence in rabbit embryos. Related to Figure 1. (A)** Epifluoresence imaging and immunostaining of GFP in uninjected rabbit embryos (left panel) and in rabbit embryos injected with primed rhesus PSCs not expressing GFP (n = 40). **(B)** Immunostaining of GFP in primed LyonES-tGFP-(S3) rhesus PSCs. **(C)** Immunostaining of GFP in a late blastocyst-stage (E5) rabbit embryo after microinjection of rabbit naïve PSCs into morula-stage (E2) embryos (confocal imaging; scale bars: 50 $\mu$m).

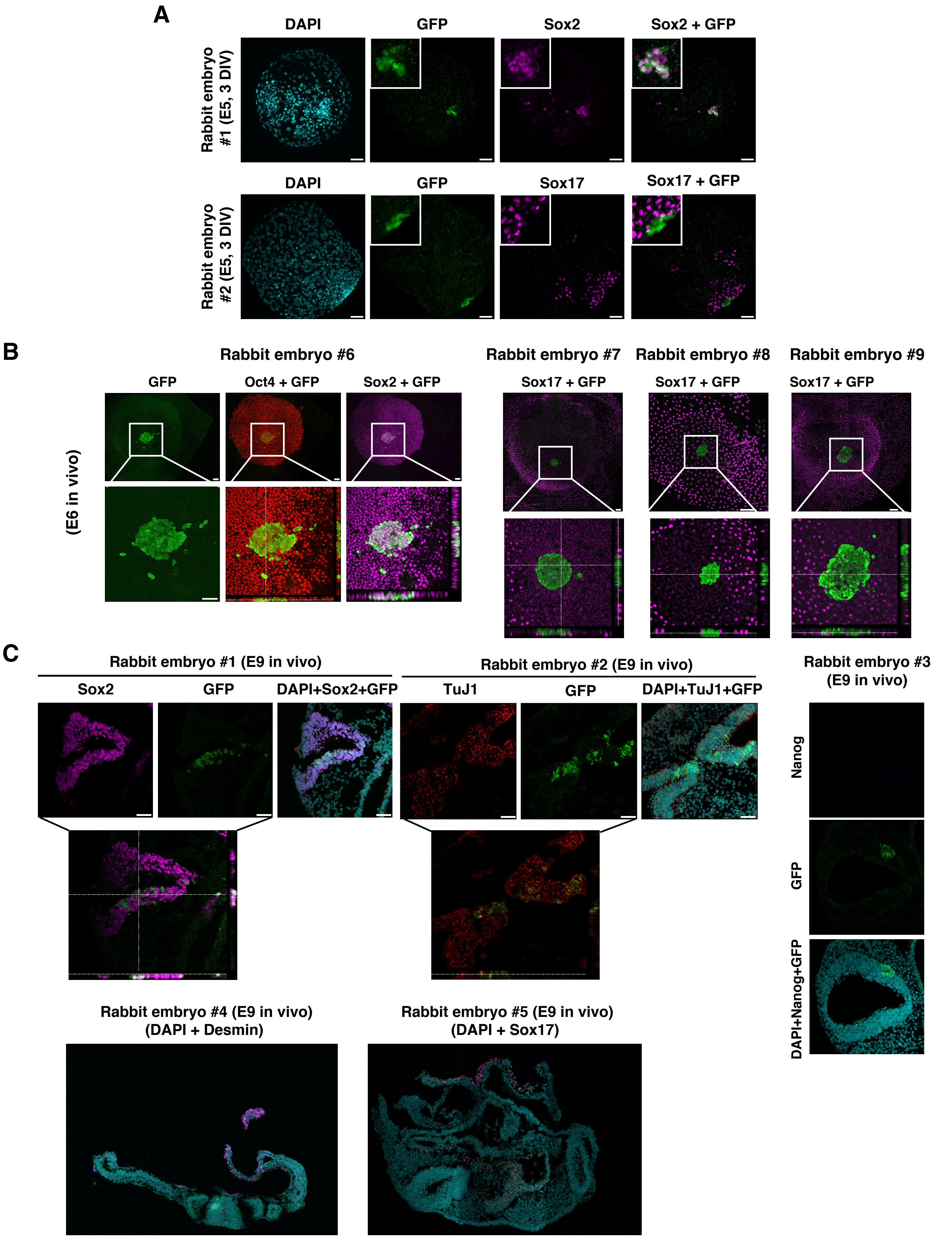

**Figure S3: Colonization of rabbit embryos by mouse ESCs. Related to Figure 1. (A)** Immunostaining of GFP, Sox2 and Sox17 in rabbit embryos at the late blastocyst-stage (E5) after microinjection of mESC-GFP-2i/LIF cells into morula-stage embryos (confocal imaging; scale bars: 50 $\mu$m) (n = 29). **(B)** Immunostaining of GFP, Oct4, Sox2 and SOX17 performed on rabbit embryos at the pre-gastrula stage (E6) after microinjection of mESC-GFP cells into morula-stage (E2) embryos and transfer to a surrogate mother. Bottom panels represent orthogonal sections. (confocal imaging; scale bars: 50 $\mu$m). **(C)** Immunostaining of GFP, Sox2, TuJ1, Nanog, Desmin, and Sox17, in rabbit embryos at the post-implantation stage (E9) after microinjection of mESC-GFP cells into morula-stage embryos (confocal imaging; scale bars: 50 $\mu$m).

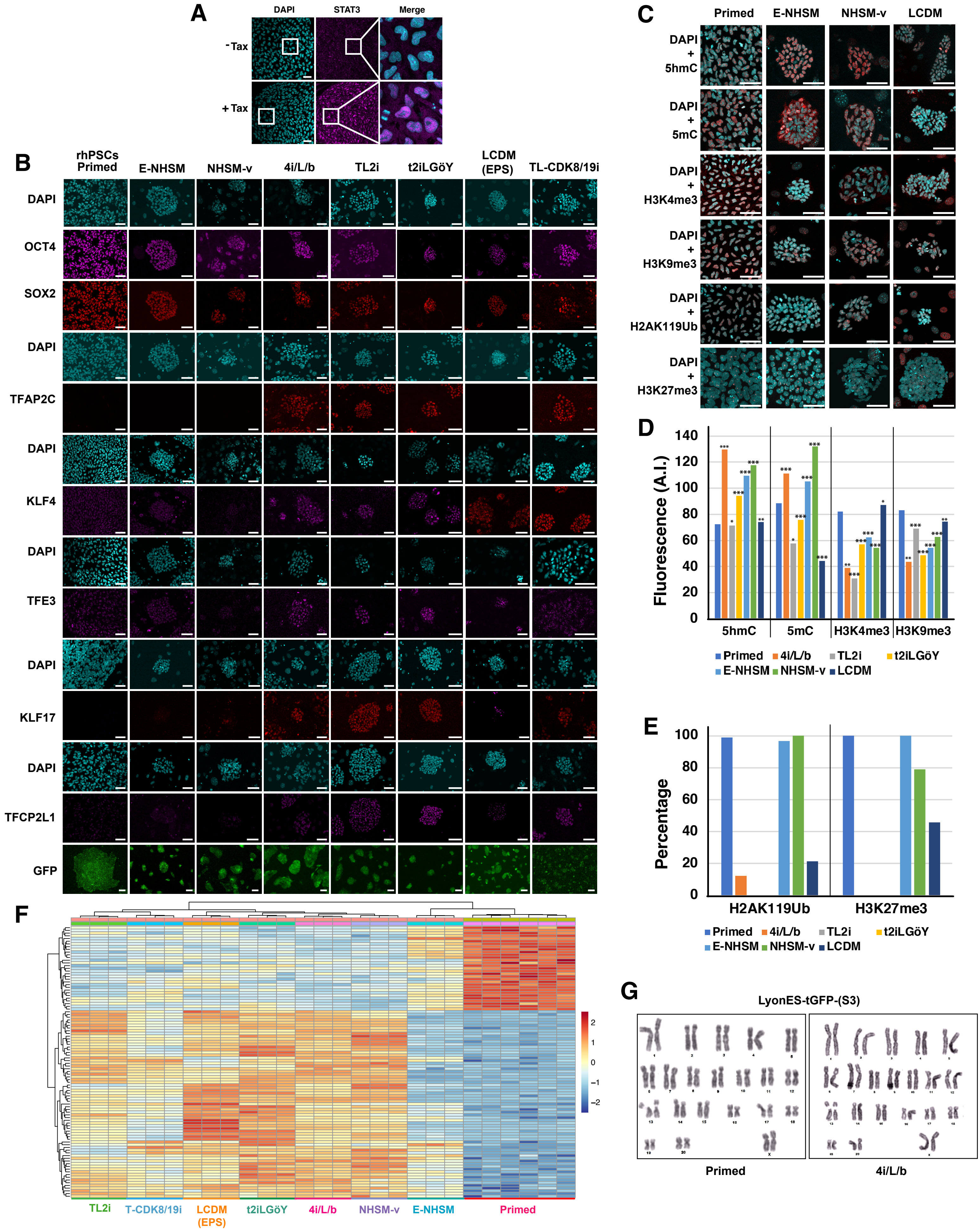

**Figure S4: Characterization of LyonES-tGFP-(S3) cells. Related to Figure 2.** **(A)** Immunolabeling of STAT3 in LyonES-tGFP-(S3) cells, before and after treatment with 250nM tamoxifen for 48 h (confocal imaging; scale bars: 50 $\mu$m). **(B) (C)** Immunolabeling of pluripotency markers and methylation marks in LyonES-tGFP-(S3) cells, before and after applying culture protocols for naïve conversion (confocal imaging; scale bars: 50 $\mu$m). **(D)** Histogram representing the quantification of 5hmC, 5mC, H3K4me3 and H3K9me3 methylation marks (Student's t test, p-value ≤ 0,05 : *; p-value ≤ 0,01 : **; p < 0,001: ***). **(E)** Histogram representing the percentage of H2AK119Ub and H3K27me3 marks. **(F)** Heatmap of transcriptome data (mean values/cell category most differentially expressed 100 genes, listed in Table S2) using Pearson correlation coefficient as a measure of Euclidian distance between rows and between columns. (G) G-banding karyotype of LyonES-tGFP-(S3) primed cells (left panel) and a batch of LyonES-tGFP-(S3) naïve 4i/L/b cells.

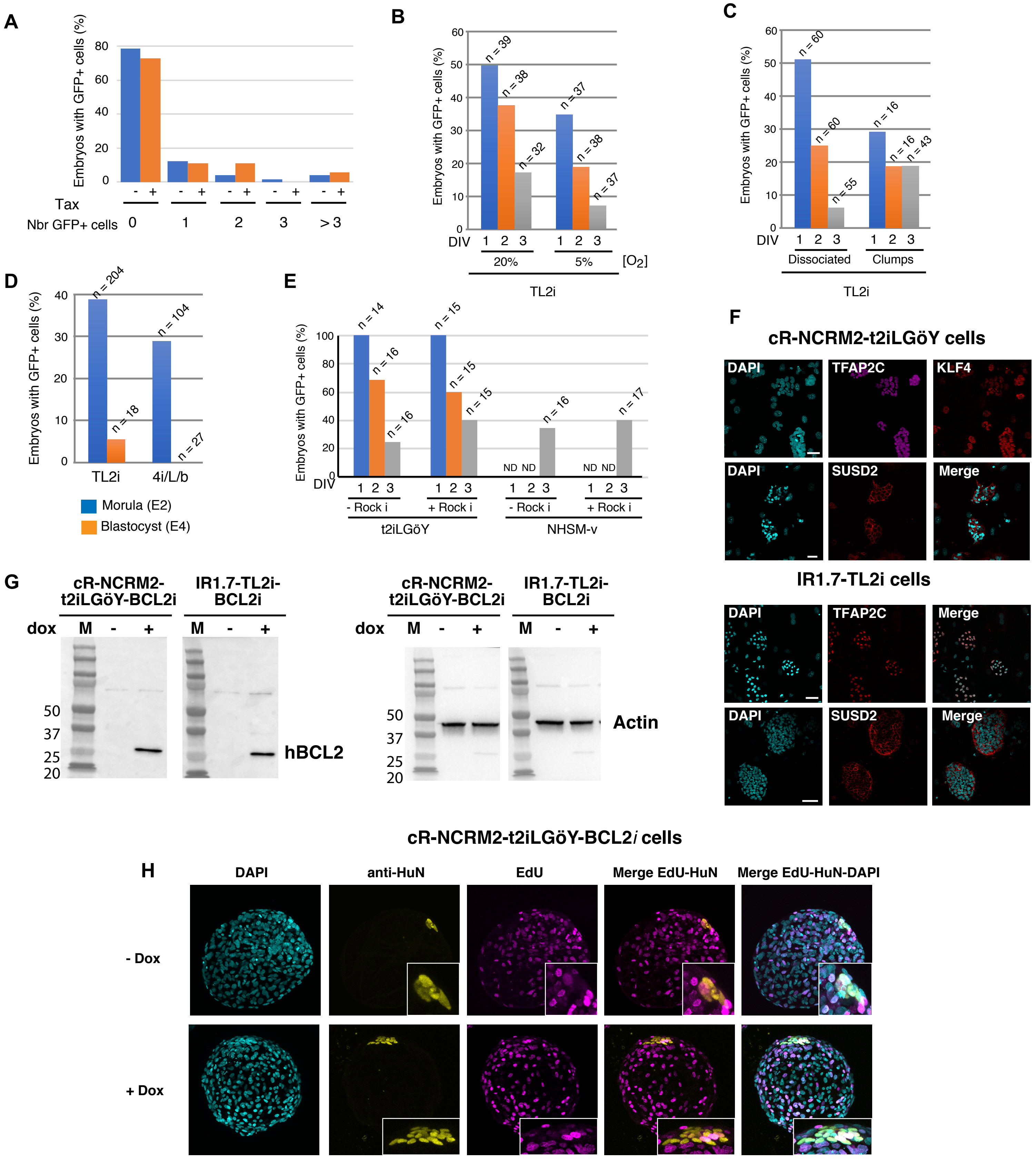

**Figure S9: Optimization of the conditions for microinjection and Colonization of rabbit embryos by human PSCs. Related to Figures 3 and 4. (A)** Histogram of percentage of rabbit embryos with GFP+ cells after microinjection of rhesus TL2i cells at the morula-stage (E2) and in vitro culture for 3 days in gradual medium in the presence (+) or in the absence (-) of 250nM tamoxifen. **(B)** Histogram of percentage of rabbit embryos with GFP+ cells after microinjection of rhesus TL2i cells at the morula-stage (E2) under normoxic (20%) and hypoxic (5%) conditions. Analyses were performed at 1, 2, and 3 DIV (n = 221). **(C)** Histogram of the percentage of rabbit embryos with GFP+ cells after microinjection of dissociated or clumps of rhesus TL2i cells into morula-stage (E2) embryos. Analyses were performed at 1, 2, and 3 DIV (n = 250). **(D)** Histogram of the percentage of rabbit embryos with GFP+ cells after microinjection of rhesus 4i/L/b and TL2i cells into morula-stage (E2) or blastocyst-stage (E4) embryos. Analyses were performed at 3 DIV (n = 353). **(E)** Histogram of the percentage of rabbit embryos with GFP+ cells after microinjection of rhesus t2iLGöY and E-NHSM cells into morula-stage (E2) embryos, in the presence or absence of ROCK inhibitor (n = 124). Analyses were performed at 3 DIV. **(F)** Immunostaining of TFAP2C and SUSD2 in NCRM2-t2iLGöY and IR1.7 TL2i cells (confocal imaging; scale bars: 50 $\mu$m). **(G)** Western blots of hBCL2 in cR-NCRM2- t2iLGöY and IR1.7-TL2i cells before (-) and after (+) treatment with 1 $\mu$g/mL doxycycline for 72 hours. **(H)** Immunostaining of HuN anf EdU in late blastocyst-stage rabbit embryos (E5, 3 DIV) after microinjection of human cR-NCRM2-t2iLGöY-BCL2*i* PSCs before (-Dox) and after (+Dox) treatment with 1 $\mu$g/mL doxycycline from 24 hours before injection to 72 hours after injection.

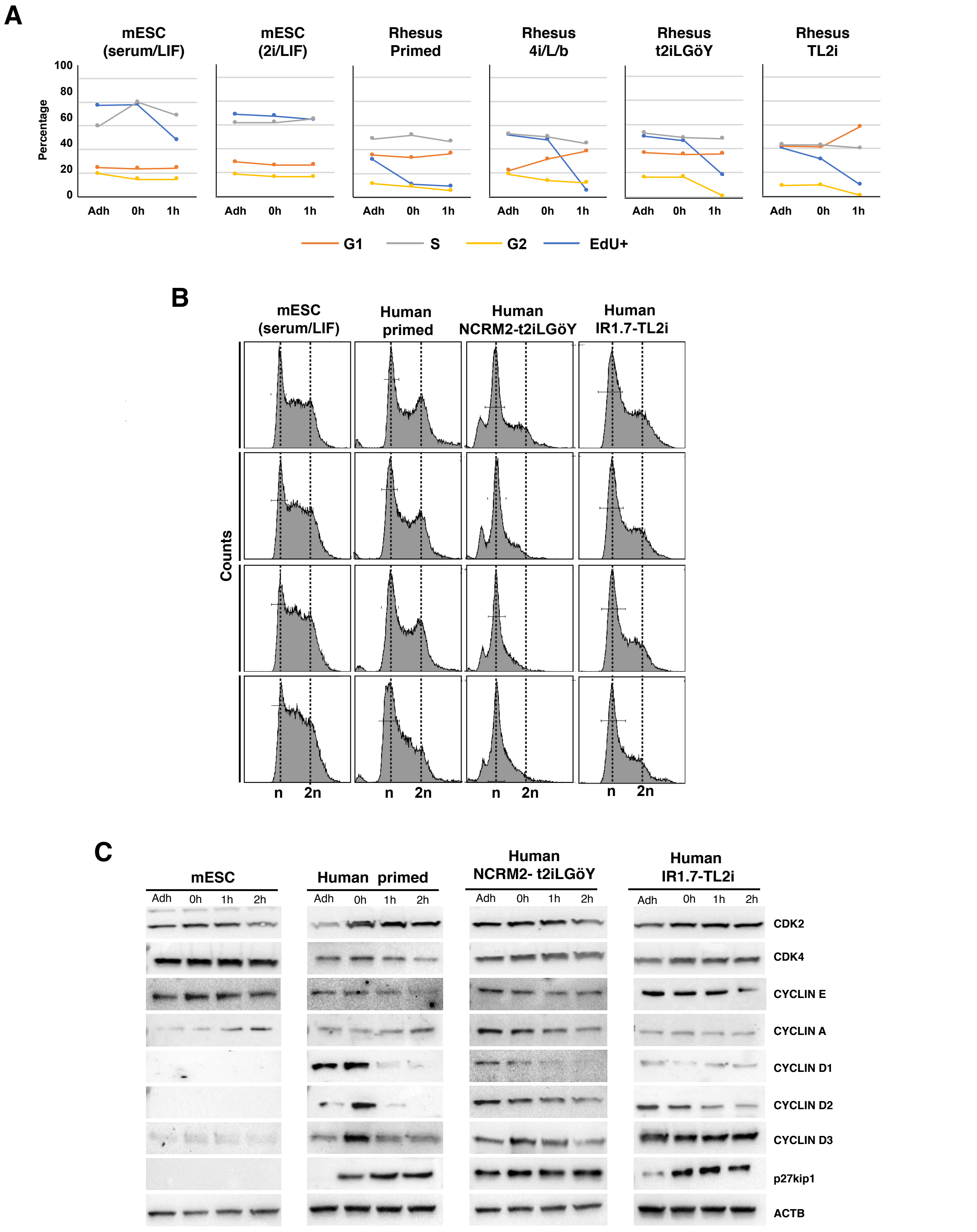

**Figure S10: Cell-cycle parameters of mouse ESCs, rhesus PSCs, and human iPSCs, before and after reprogramming to naïve states. Related to Figure 6. (A)** Histograms showing the cell cycle distribution of mESCs (serum/LIF and 2i/LIF) and rhesus PSCs (primed, 4i/L/b, t2iLGöY and TL2i) in “Adh”, “0h”, “1h”, and “2h” conditions analyzed by flow cytometry after EdU incorporation and PI staining (% EdU+, G1, and S/G2 cells). **(B)** Cell cycle distribution of mESCs (serum/LIF) and human PSCs (primed, cR-NCRM2-t2iLGöY and IR7.1-TL2i) in “Adh”, “0h”, “1h”, and “2h” conditions analysed by flow cytometry after PI staining. **(C)** Western blot analysis of cell-cycle regulators in mESCs (serum/LIF) and human iPSCs (primed, cR-NCRM2-t2iLGöY and IR7.1-TL2i) in “Adh”, “0h”, “1h”, and “2h” conditions (results of 3 experiments).

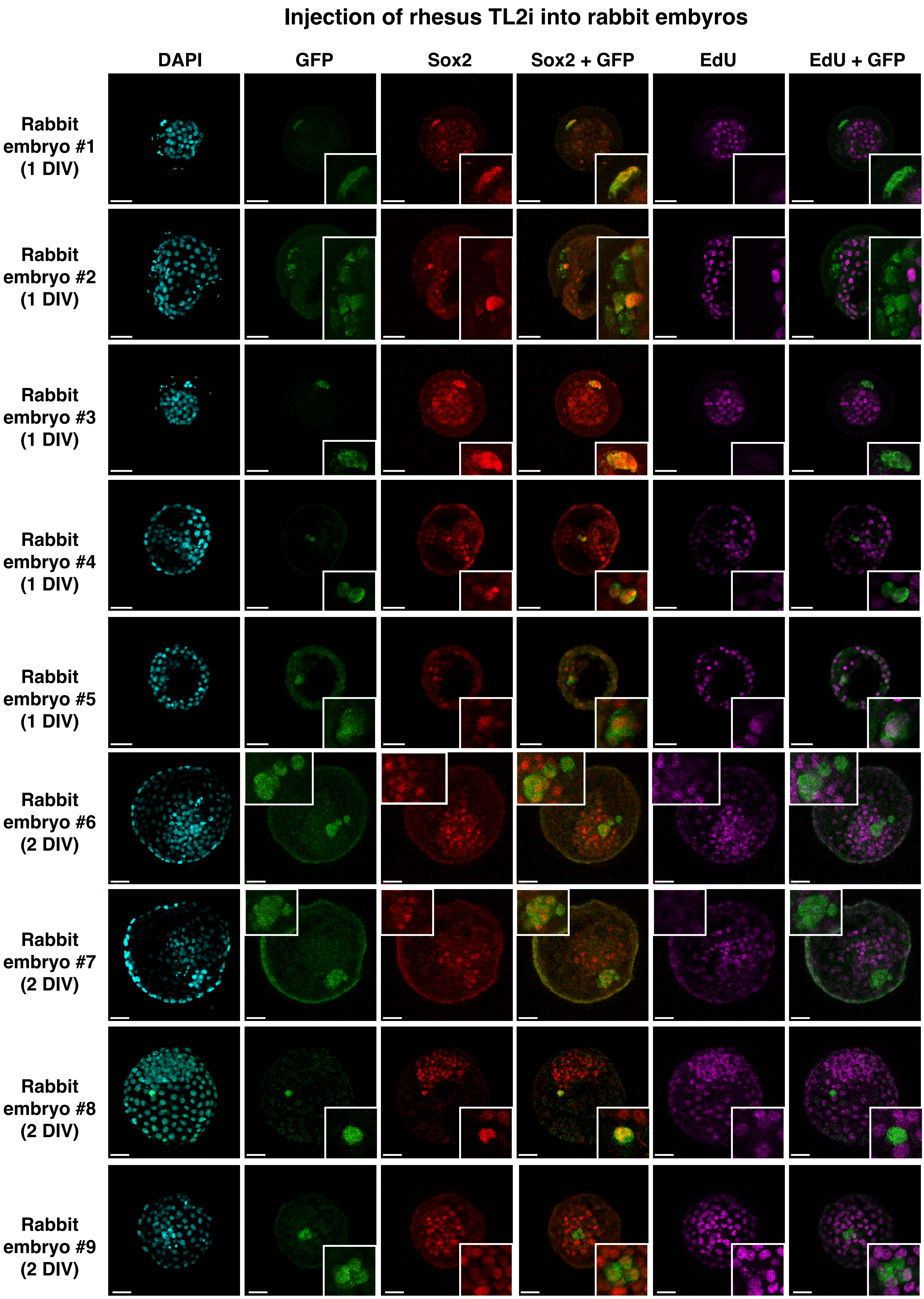

**Figure S11: DNA replication of Rhesus PSCs after microinjection into rabbit embryos. Related to Figure 7.** Immunostaining of GFP, Sox2, and EdU in late blastocyst-stage rabbit embryos (E5, 3 DIV) after microinjection of rhesus TL2i cells into morula-stage (E2) embryos (confocal imaging; scale bars: 50 $\mu$m).

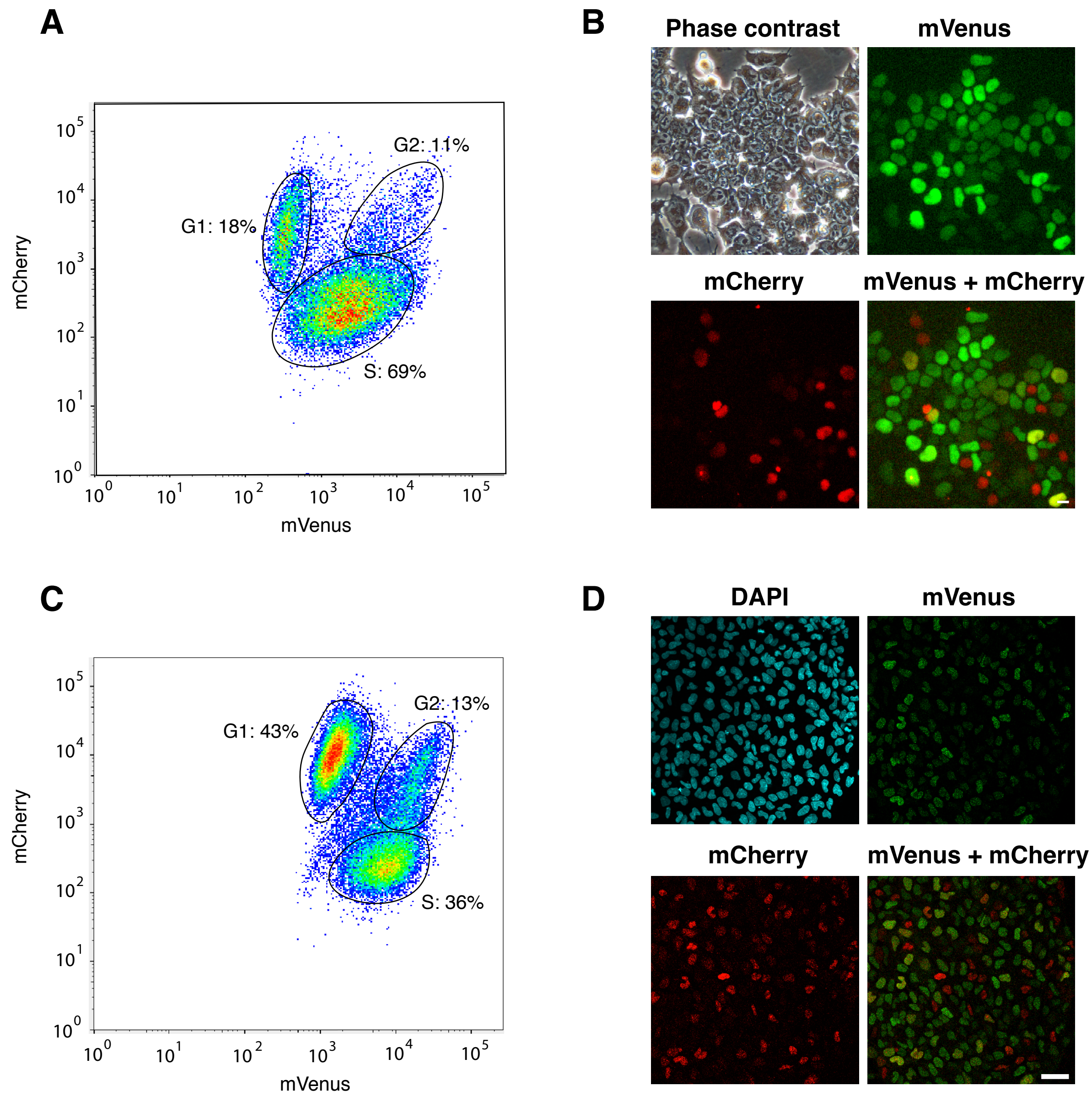

**Figure S12: Characterization of mouse ESCs and rhesus PSCs harboring the FUCCI(CA) cell cycle reporter. Related to Figure 7. (A)** Flow cytometry analysis and **(B)** epifluorescence imaging of mESCs-FUCCI (CA) cells. **(C)** Flow cytometry analysis and **(D)** epifluorescence imaging of rhesus primed-FUCCI (CA) cells (scale bars: 50 $\mu$m).

### Supplemental Experimental Procedures

#### Ethics

All procedures in macaque monkeys and rabbits were approved by the French ethics committee CELYNE (approval number: APAFIS#4858 and APAFIS#6438). Chimera experiments involving human iPSCs were approved by the INSERM ethics committee.

#### Cell lines, media composition, and culture

mESC lines were routinely cultured in Glasgow's modified Eagle's medium (Gibco) supplemented with 10% fetal calf serum (CRC0406; PerbioScience) and 1,000 U/mL LIF on gelatin-coated dishes. They were also cultured in N2B27 supplemented with 1,000 U/mL LIF, 1 μM PD0325901, and 3 μM CHIR99021 (2i/LIF), as indicated. Both the LyonES-tGFP and Lyon-ES-tGFP-(S3) rhesus PSC lines (Wianny et al., 2008, Chen et al., 2015a) and the human iPSC line IR7.1 (Chen et al., 2015a), (Ng et al., 2012) cell lines were routinely cultured at 37 °C in 5% $CO_2$ and 5% $O_2$ in knockout Dulbecco's modified Eagle's medium (KO-DMEM) supplemented with 20% knockout serum replacement (KOSR), 1 mM glutamine, 0.1 mM β-mercaptoethanol (Sigma), 1% non-essential amino acid (Gibco), and 4–8 ng/mL FGF2 (Gibco) on growth-inactivated murine embryonic fibroblasts.

Primed to naïve conversion was performed using previously described protocols, including E-NHSM (https://hannalabweb.weizmann.ac.il) (Gafni et al., 2013), NHSM-v (Chen et al., 2015b), 4i/L/b (Fang et al., 2014), 5iL/A (Theunissen et al., 2014), t2iLGoY (Guo et al., 2017), TL2i (Chen et al., 2015a), TL-CDK8/19i [modified from (Lynch et al., 2020)], and LCDM (EPS) (Yang et al., 2017a). Primed cells were dissociated and plated on fresh feeder cells for 24 h before shifting to the naïve culture media. Fresh medium was added daily to the culture plates. The medium composition for E-NHSM reprogramming (Gafni et al., 2013) was as follows: Neurobasal: DMEM-F12 (1:1) supplemented with N2B27 (Gibco), L-ascorbic acid, 0.5% KOSR, 10 ng/mL human LIF (Pepeotech), 5 μM IWR1 (Sigma), 1.5 μM CHIR99021 (Miltenyi Biotec), 1 μM PD0325901 (Miltenyi Biotec), 2 μM BIRB796 (Axon), 5 μM SP600125 (Tocris), 2 μM Gö6983 (Tocris), 20 ng/mL activin A (Peprotech), and 1 μM CGP77675 (Axon). Cells were passaged every 3–4 days by single-cell dissociation using Accutase (Gibco). ROCK inhibitor (Y-27632; Miltenyi Biotec) was added during passaging for 24 h. The NHSM-v protocol is a modified version of the NHSM protocol applied to cynomolgus macaque ESCs (Chen et al., 2015b). These cells were cultured in KO-DMEM supplemented with 20% KOSR, 4 ng/mL FGF2 (Gibco), 10 ng/mL hLIF (Peprotech), 3 μM CHIR99021 (Miltenyi Biotec), 0.5 μM PD0325901 (Miltenyi Biotec), 5μM SP600125 (Tocris), and 10 μM SB203580 (Tocris). Cells were passaged every 3–4 days by single-cell dissociation using 1 mg/mL Accutase (Gibco). ROCK inhibitor (Y-27632; Miltenyi Biotec) was added during passaging for 24 h. The 4i/L/b media was adapted from the protocol published by Fang *et al.* (Fang et al., 2014). Cells were cultured in KO-DMEM supplemented with 20% KOSR (Gibco), 0.5 μM PD0325901 (Miltenyi Biotec), 3 μM CHIR99021 (Miltenyi Biotec), 10 μM SP600125 (Tocris), 100 μM SB203580 (Tocris), 10 ng/mL hLIF (Peprotech), and 2.5 ng/mL FGF2 (Gibco). Cells were passaged every 3–4 days by single-cell dissociation using 0.05% Trypsin-ethylenediaminetetraacetic acid (Trypsin-EDTA; Gibco). ROCK inhibitor (Y-27632; Miltenyi Biotec) was added during passaging for 24 h. Primed cells were converted to 5i/LA state using the following medium composition (Theunissen et al., 2014): DMEM-F12 (1:1) medium supplemented with N2 (Gibco), B27 (Gibco), 0.5% KOSR (Gibco), 1 μM PD0325901 (Miltenyi Biotec), 1 μM IM-12 (Enzo), 0.5 μM SB590885 (Tocris), 1 μM WH-4-023 (Tocris), and 20 ng/mL activin A (Peprotech). ROCK inhibitor (Y-27632; Miltenyi Biotec) was added during passaging for 24 h. Primed cells were converted to the t2iLGöY state using the following medium composition (Guo et al., 2017): first in cRM1 media for three days with 1 μM PD0325901 (Miltenyi Biotec), 10 ng/mL hLIF (Peprotech) and 1mM Valproic acid (VPA), followed by two days in cRM2 media with DMEM-F12 (1:1) medium supplemented with N2, B27 (Gibco), 1 μM PD0325901 (Miltenyi Biotec) and 250 μM Vitamin C (Sigma-Aldrich) and then the final media containing DMEM-F12 (1:1) medium supplemented with N2, B27 (Gibco), 1 μM PD0325901 (Miltenyi Biotec), 1,000 U/mL LIF, 1 μM Gö6983 (Bio-techne), 2 μM XAV939 (Sigma). Cells were passaged every 4–5 days by single-cell dissociation using Tryple (Gibco). ROCK inhibitor (Y-27632; Miltenyi Biotec) was added during passaging for 24 h. For reprogramming to the TL2i state, PSCs expressing the inducible fusion protein STAT3-$ER^{T2}$ were cultivated in the TL2i medium composed of KO-DMEM (Gibco), 20% KOSR (Gibco), 10,000 U/mL LIF, 3 μM CHIR99021 (Miltenyi Biotec), 1 μM PD0325901 (Miltenyi Biotec), and 250 nM 4′-Hydroxy-Tamoxifen (4′OHT; Calbiochem). Cells were passaged every 3–4 days by single-cell dissociation using 0.05% Trypsin-EDTA (Gibco). ROCK inhibitor (Y-27632; Miltenyi Biotec) was added during passaging for 24 h. For TL-CDK8/19i protocol, cells were cultured in medium composed of KO-DMEM (Gibco), 20% KOSR (Gibco), 10,000 U/mL LIF, and 1 μM CNIO-47799. Cells were passaged every 3–4 days by single-cell dissociation using 0.05% Trypsin-EDTA (Gibco). ROCK inhibitor (Y-27632; Miltenyi Biotec) was added during passaging for 24 h.Primed cells were converted to the LCDM (EPS) (Yang et al., 2017) state using medium with the following composition: DMEM-F12 (1:1), N2 (Gibco), B27 (Gibco), 5% KOSR (Gibco), 10 ng/mL hLIF (Peprotech), 1 μM CHIR99021 (Miltenyi Biotec), 2 μM (S)-(+)-Dimethindene maleate (Tocris), 2 μM minocycline hydrochloride (Santa Cruz Biotechnology), and 0.75 μM IWR1 (Sellekchem). Cells were passaged every 3–4 days by single-cell dissociation using 0.05% Trypsin-EDTA (Gibco). ROCK inhibitor (Y-27632; Miltenyi Biotec) was added during passaging for 24 h (**Table S3**).

To avoid the accumulation of karyotypic abnormalities, new naïve cell batches were regularly produced and none of the naïve cells were cultured for more than 2 months after reprogramming.

**G-banding karyotyping**

G-banding karyotyping was performed by Cell Guidance Systems (www.cellgs.com).

**Generation of FUCCI reporter rhesus PSCs and mESCs**

The *PB-Puro$^R$-CAG-mVenus : hGeminin-IRES-mCherry : hCdt1* plasmid was generated as follows. First, an EcoRI/XbaI DNA fragment from the *mCherry-hCdt1(1/100)Cy(-)/pcDNA3* plasmid (Sakaue-Sawano et al., 2017) containing the mCherry : hCdt1 cassette was sub-cloned between the EcoRI and XbaI restriction sites in *PB-CAG-cHA-IRES-hyg* to generate *PB-Hygro-mcherry : hCdt1*. Second, a AlfII/XbaI fragment from the *mVenus-hGeminin(1/110)/pcDNA3* plasmid (Sakaue-Sawano et al., 2017) containing the mVenus : hGeminin cassette was subcloned between the SapI and AgeI restriction sites in *PB-Hygro-mcherry : hCdt1* to generate *PB-Hygro-mcherry : hCdt1-mVenus : hGeminin*. Third, a SapI/AgeI fragment containing the *Puro$^R$* gene and the CAG promoter was sub-cloned between the SapI and AgeI restriction sites in *PB-Hygro-mcherry : hCdt1-mVenus : hGeminin* to generate the *PB-Puro$^R$-CAG-mVenus : hGeminin-IRES-mCherry : hCdt1* plasmid. Rhesus PSCs and mESCs were transfected using the NEON transfection system according to the instructions provided by the manufacturer. Briefly, cells were dissociated and resuspended at a density of $10.10^6$ cells/mL. For transfection, 100 µL of the cell suspension was mixed with 2.5 µg of the *PB-Puro$^R$-CAG-mVenus : hGeminin-IRES-mCherry : hCdt1* vector and 2.5 µg of *PBase* plasmid. Transfection parameters used for rhesus PSCs were 1,050 V, 20 ms, and 2 pulses. Cells were plated on growth-inactivated murine embryonic fibroblasts in medium supplemented with 10 µM ROCK inhibitor (Y-27632; Miltenyi Biotec) and selected in 250 ng/mL G418. For mESCs, the parameters used were 1,200 V, 20 ms, and 2 pulses. Cells were analyzed using a FACS LSR II (Becton-Dickinson) equipped with 355, 488, and 561 nm lasers. Data were acquired and analyzed using the DiVa software.

**Generation of BCL2 inducible human PSCs**

The *BCL2* plasmid was generated as follows. First, a DNA fragment from the *pCS2+Flag-BCL2* plasmid was amplified by PCR using primers flanked by *Kpn*I and *Spe*I restriction sites at the 5' and 3' end, respectively. The PCR fragment was then subcloned in pXlone-GFP-Neo3R-P2A-TRE-EF1 plasmid. This plasmid was obtained after modification of the *Xlone-GFP* plasmid (a gift from Xiaojun Lian, Addgene plasmid #96930) in which the Blasticidin resistance gene was replaced by Neomycin. cR-NCRM2-t2iLGöY and IR1.7-TL2i cells were transfected using the NEON transfection system according to the instructions provided by the manufacturer. Briefly, cells were dissociated and resuspended at a density of $10.10^6$ cells/mL. For transfection, 100 µL of the cell suspension was mixed with 2.5 µg of the *pXlone-Flag-BCL2-Neo3R-P2A-TRE-EF1* vector and 2.5 µg of *PBase* plasmid. Transfection parameters used were 1,050 V, 20 ms, and 2 pulses. Cells were plated on growth-inactivated murine embryonic fibroblasts in medium supplemented with 10 µM ROCK inhibitor (Y-27632; Miltenyi Biotec) and selected in 250 ng/mL G418.

**Generation of rhesus and cynomolgus monkey embryos and cell microinjection**

Macaque cynomolgus embryos were produced through ovarian stimulation, followed by intra-cytoplasmic sperm injection (ICSI) of oocytes, as previously described (Tachibana et al., 2012). Briefly, females received twice-daily injections of 25 IU of recombinant human follicle-stimulating hormone (MSD laboratories) from days 1 to 8 (starting at day 1–4 of the menstrual cycle). They received 30 IU of recombinant human luteinizing hormone (Merck) on days 7 and 8. Females received an injection of gonadotropin-releasing hormone antagonist (MSD laboratories) (0.075 mg/kg body weight) on day 8, and an injection of 1,040 IU of human chorionic gonadotropin (Merck) at day 8. Estradiol, luteinizing hormone, progesterone dosage, and ultrasonographic scans were performed to monitor ovarian response. Follicle aspiration and oocyte retrieval were performed by laparoscopy 36 h after injection of human chorionic gonadotropin. Oocytes were transferred to HEPES-buffered TALP medium containing 3 mg/mL of bovine serum albumin (Sigma). Cumulus and granulosa cells were removed by incubating the oocytes for 30 s with hyaluronidase (300 µg/mL). Metaphase II stage oocytes were selected for ICSI and transferred to drops of hamster embryo culture medium-9 (HECM9) covered by liquid paraffin (Origio) at 37 °C in a 5% $O_2$ + 5% $CO_2$ atmosphere. Four days after fertilization (E4), morula-stage embryos were microinjected with 10 cells and further cultured in drops of PSC medium for 4 h. PSC media were those used for culturing the cells prior to injection, depending on the reprogramming protocols. After 4 h, embryos were transferred into a 1:1 mix of PSC medium and HECM9 and further cultured for 20 h. After 24 h, embryos were transferred to HECM9 and further cultured.

**Production of rabbit embryos, cell microinjection, embryo culture, and transfer**

Rabbit embryos were produced by ovarian stimulation. Briefly, sexually mature New Zealand white rabbits were injected with follicle-stimulating hormone and gonadotropin-releasing hormone, followed by artificial insemination or breeding, as previously described. Eight-cell-stage embryos (E1.5) were flushed from explanted oviducts 36–40 h after insemination and cultured in a 1:1:1 mixture of RPMI 1640 medium, DMEM, and Ham's F10 (RDH medium; Thermo Fisher Scientific) at 38 °C in 5% $CO_2$ until cell microinjection. For the latter procedure, 10 cells were microinjected under the zona pellucida of morula (E2.5)- or blastocyst (E4)-stage rabbit embryos. The embryos were further cultured using the same experimental procedure as for monkey embryo. For embryo transfer, surrogate mothers were prepared through intramuscular injection of 1.6 μg of buserelin acetate (Intervet). Morula-stage embryos (6–8) were transferred to each oviduct of the recipient by laparoscopy. Four days after transfer, pre-implantation embryos (E6) were recovered by flushing the explanted uterine horns. Post-implantation-stage embryos (E9) were recovered by dissecting the uterine horns.

**Immunoblotting**

For immunoblotting, cells were grown on matrigel to avoid contamination by mouse embryonic fibroblasts. Frozen cell pellets were lysed in RIPA buffer complemented with protease and phosphatase inhibitors. Protein lysates were then cleared by centrifugation (14,000 r.p.m. for 30 min). After SDS–PAGE and electroblotting on polyvinylidene difluoride, the membranes were incubated with specific primary antibodies (antibodies used in this study are listed below). Blots were incubated with horseradish peroxidase-coupled anti-mouse or -rabbit immunoglobulin G (Jackson ImmunoResearch) and developed with Clarity Western ECL Substrate (BIO-RAD).

**Histology, histochemistry, immunostaining**

Monkey (E7) and rabbit (E3, E4, E5, and E6) pre-implantation embryos were fixed in 2% paraformaldehyde (PFA) for 20 min at room temperature. After three washes in phosphate-buffered saline (PBS) containing 0.1% Tween-20 (PBS-0.1%T), embryos were permeabilized in PBS-1%T overnight at 4 °C on a rotating shaker. Embryos were subsequently placed in blocking solution (PBS-0.1%T) supplemented with 5% donkey serum for 1 h at room temperature. There were incubated with primary antibodies diluted in blocking solution overnight at 4°C (antibodies used in this study are listed in table S4). After four washes (3 × 5 min + 1 × 30 min) in PBS-0.1%T, embryos were incubated in secondary antibodies diluted in blocking solution at a concentration of 1:300 for 1 h at room temperature and transferred through several washes of PBS-0.1%T before staining the nuclei with 4′,6-diamidino-2-phenylindole (DAPI; 0.5 μg/mL). Embryos were analyzed by confocal imaging (DM 6000 CS SP5; Leica). Acquisitions were performed using a water immersion objective (25×/1.25 0.75, PL APO HCX; Leica).

E8 rabbit post-implantation embryos were fixed in 2% PFA at 4 °C for 1 h. For cryoprotection, embryos were placed in 10% sucrose for 1 h followed by 30% sucrose overnight at 4 °C. Embryos were embedded in NEG50 compound (Invitrogen) and frozen at −80 °C. Sections of 15–20 μm were generated using a Microm HM550 cryostat and maintained at −80 °C until immunostaining was performed. For immunostaining, sections were thawed for 30 min at room temperature, saturated with PBS for 15 min at room temperature, and permeabilized (as necessary) with three baths of PBS containing 0.5% Triton X100 (Sigma-Aldrich). Sections were incubated in blocking solution: PBS with 0.1% Triton X100 and 10% donkey serum (Jackson Immunoresearch Laboratory) for 30 min. Subsequently, the sections were incubated with primary antibodies overnight at 4 °C. After several washes, the sections were incubated with the appropriate fluorescence-conjugated secondary antibodies at room temperature for 1 h. Nuclei were stained with DAPI (0.5 μg/mL). Sections were mounted on coverslips with Fluoromount G (Thermo Fisher Scientific). Tiled scans were automatically acquired using the LAS AF software (Leica).

Cells were fixed with 2% PFA in PBS at room temperature for 20 min, washed thrice (10 min each) with PBS and permeabilized in PBS containing 0.5% Triton X100. Non-specific binding sites were blocked using PBS supplemented with 10% donkey serum for 1 h at room temperature. The cells were incubated overnight at 4 °C with primary antibodies. After three rinses (10 min each) with PBS, the cells were incubated with secondary antibodies at room temperature for 1 h. The nuclei were stained with DAPI (0.5 μg/mL). The cells were mounted on coverslips by using the mounting medium M1289 (Sigma). Cells and embryo sections were analyzed by confocal imaging (DM 6000 CS SP5; Leica). Acquisitions were performed using an oil immersion objective (45×/1.25 0.75, PL APO HCX; Leica).

EdU staining was performed using the Click-iT™ Edu Imaging kit (Fisher Scientific). Briefly, embryos microinjected with PSCs were incubated for 1 h with 10 μM EdU in RDH medium. Embryos were fixed with 2–4% PFA, and immunostaining was performed according to the instructions provided by the manufacturer. Embryos were analyzed by confocal imaging (DM 6000 CS SP5; Leica), and acquisitions were performed using a water immersion objective (25×/1.25 0.75, PL APO HCX; Leica).

**Two-photon recording**

Embryos microinjected with rhesus and mESCs were placed in drops (5 µL) of medium and covered with liquid paraffin (Origio). For recording, embryos were cultured under the inverted Axio-Observer Z1 (Zeiss) two-photon microscope for 3 days, until they reach the E5 stage. Images were acquired using the Zeiss Zen software, and image analysis was performed using the ImageJ software.

**Flow cytometry analysis of cell cycle distribution**
Cells were labeled for 15 minutes with 10µM EdU in the culture medium using the Click-it Plus EdU flow cytometry kit (ThermoFischer). Cells were then washed and processed for the detection of incorporated EdU according to the manufacturer's instructions. After immunostaining, cells were incubated for 20 min with 1 mg/ml RNAse in PBS-0.13 mM EGTA. Propidium iodide (1 □g/ml) was added just before analysis. Cells were analyzed using a FACS LSR II (Becton-Dickinson) equipped with 355, 488, and 561 nM lasers. Data were acquired and analyzed using DiVa software.

**RNA sequencing**
RNA from cell lines was extracted from 4–5.$10^6$ cells using the RNeasy mini-kit (Qiagen). The libraries were prepared using 200 ng of RNA with the NextFlex Rapid Directional mRNA-Seq kit (Bioo-Scientific). Samples were sequenced on a NextSeq500 sequencing machine (Illumina) as single reads of 75 bp. The bcl2fastq conversion software was used for demultiplexing (Illumina), and data trimming was performed using Cutadapt. The sequencing depth for each sample was approximately 30 million reads, which were subsequently mapped to the Mmul8 rhesus macaque genome using the HISAT2 alignment program and quantified with the HTseq script.

**Bioinformatics analysis**
RNA-seq data obtained from rhesus primed and naïve cell lines were analyzed using the R software. PCA, differential expression analysis (with false discovery rate <0.1), and hierarchical clustering were performed with DESeq2 (Love et al., 2014). For cell lines, PCA was performed using the top 500 genes selected by highest row variance. For the comparison of cell lines with embryo data, we used the Seurat R package (version 3.1). Briefly, the two compendia of datasets were transformed to a Seurat object and merged. The transcript counts were subsequently log-transformed and normalized. Finally, the variable genes (3,000) were used as input for PCA.

**Table S1:** Summary of interspecies chimera experiments. Related to Figure 1, 2, 4 and 5.

| | Rabbit host | | |
|---|---|---|---|
| | E3 (1 DIV) | E4 (2 DIV) | E5 (3 DIV) |
| Rhesus PSCs | | | |
| Primed | | | |
| **Nb GFP+ embryos/injected embryos** | **10/104** | **0/89** | **0/90** |
| Nb NANOG+ embryos/GFP+ embryos | 0/7 | | |
| Nb SOX2+ embryos/GFP+ embryos | 0/2 | | |
| Nb GATA6+ embryos/GFP+ embryos | 0/1 | | |
| TL2i | | | |
| **Nb GFP+ embryos/injected embryos** | **109/141** | **78/145** | **79/204** |
| Nb NANOG+ embryos/GFP+ embryos | 14/16 | 7/12 | 2/7 |
| Nb SOX2+ embryos/GFP+ embryos | 45/53 | 30/52 | 8/40 |
| Nb GATA6+ embryos/GFP+ embryos | 6/31 | 8/22 | 14/33 |
| 4i/L/b | | | |
| **Nb GFP+ embryos/injected embryos** | **45/59** | **32/65** | **30/104** |
| Nb NANOG+ embryos/GFP+ embryos | 5/7 | 3/12 | 1/7 |
| Nb SOX2+ embryos/GFP+ embryos | 3/5 | 0/1 | 1/6 |
| Nb GATA6+ embryos/GFP+ embryos | 4/33 | 5/19 | 6/17 |
| t2iLGoY | | | |
| **Nb GFP+ embryos/injected embryos** | **92/96** | **72/97** | **69/114** |
| Nb NANOG+ embryos/GFP+ embryos | 10/40 | 0/21 | 0/11 |
| Nb SOX2+ embryos/GFP+ embryos | 14/43 | 6/31 | 3/10 |
| Nb GATA6+ embryos/GFP+ embryos | 3/39 | 10/38 | 8/20 |
| LCDM (EPS) | | | |
| **Nb GFP+ embryos/injected embryos** | **66/89** | **36/89** | **44/157** |
| Nb NANOG+ embryos/GFP+ embryos | 0/19 | 2/12 | 10/17 |
| Nb SOX2+ embryos/GFP+ embryos | 20/31 | 11/15 | 5/20 |
| Nb GATA6+ embryos/GFP+ embryos | 1/24 | 0/13 | 2/19 |
| E_NHSM | | | |
| **Nb GFP+ embryos/injected embryos** | **15/71** | **1/70** | **0/71** |
| Nb NANOG+ embryos/GFP+ embryos | 0/24 | | |
| Nb SOX2+ embryos/GFP+ embryos | 0/20 | | |
| Nb GATA6+ embryos/GFP+ embryos | 2/27 | | |
| NHSM_v | | | |
| **Nb GFP+ embryos/injected embryos** | **11/68** | **8/84** | **10/118** |
| Nb NANOG+ embryos/GFP+ embryos | 0/3 | | |
| Nb SOX2+ embryos/GFP+ embryos | 1/3 | 0/3 | 0/7 |
| Nb GATA6+ embryos/GFP+ embryos | 0/5 | 0/5 | 1/3 |
| TL_CDK8/19ii | | | |
| **Nb GFP+ embryos/injected embryos** | **43/79** | **19/80** | **10/101** |
| Nb NANOG+ embryos/GFP+ embryos | 10/20 | 2/7 | 1/4 |
| Nb SOX2+ embryos/GFP+ embryos | 2/12 | 2/11 | 1/5 |
| Nb GATA6+ embryos/GFP+ embryos | 6/15 | 1/3 | 0/1 |
| Mouse ESCs | | | |
| Serum + LIF | | | |
| **Nb GFP+ embryos/injected embryos** | | | **235/238** |
| Nb NANOG+ embryos/GFP+ embryos | | | 77/77 |
| Nb SOX2+ embryos/GFP+ embryos | | | 58/58 |
| Nb SOX17+ embryos/GFP+ embryos | | | 0/30 |
| 2iLIF | | | |
| Nb GFP+ embryos/injected embryos | | | 19/29 |
| Nb SOX2+ embryos/GFP+ embryos | | | 7/7 |
| Nb SOX17+ embryos/GFP+ embryos | | | 0/7 |
| **Total** | **775** | **786** | **1226** |
| **Total injected embryos** | **2787** | | |

| | Rabbit host | | |
|---|---|---|---|
| | E3 (1 DIV) | E4 (2 DIV) | E5 (3 DIV) |
| Human PSCs | | | |
| hiPSCs_t2iLGoY | | | |
| **Nb GFP+ embryos/injected embryos** | **27/29** | **20/31** | **97/350** |
| Nb NANOG+ embryos/GFP+ embryos | | | 5/23 |
| Nb SOX2+ embryos/GFP+ embryos | 24/29' | 11/20 | 45/97 |
| Nb GATA6+ embryos/GFP+ embryos | | | 3/14 |
| Nb SOX17+ embryos/GFP+ embryos | | | 15/30 |
| Nb T-BRA+ embryos/GFP+ embryos | | | 4/11 |
| Nb NESTIN+ embryos/GFP+ embryos | | | 0/5 |
| hiPSCs_TL2i | | | |
| **Nb GFP+ embryos/injected embryos** | **33/39** | **18/36** | **75/298** |
| Nb NANOG+ embryos/GFP+ embryos | 15/15 | 3/7 | 0/9 |
| Nb SOX2+ embryos/GFP+ embryos | 9/11 | 4/8 | 2/33 |
| Nb GATA6+ embryos/GFP+ embryos | 1/7 | 2/3 | 8/22 |
| Nb SOX17+ embryos/GFP+ embryos | | | 26/30 |
| Nb T-BRA+ embryos/GFP+ embryos | | | 0/7 |
| Nb NESTIN+ embryos/GFP+ embryos | | | 0/5 |
| **Total** | **68** | **67** | **648** |
| **Total injected embryos** | **783** | | |

**BCL2 inducible cell lines**

| | Rabbit host | |
|---|---|---|
| | E5 (3 DIV) | |
| hiPSCs_t2iLGoY_BCL2 | | |
| | **-Dox** | **+Dox** |
| **Nb GFP+ embryos/injected embryos** | **20/66** | **82/82** |
| Nb NANOG+ embryos/GFP+ embryos | 10/26 | 15/27 |
| Nb SOX2+ embryos/GFP+ embryos | 8/20 | 0/28 |
| Nb GATA6+ embryos/GFP+ embryos | 4/20 | 4/27 |
| hiPSCs_TL2i_BCL2 | | |
| | **-Dox** | **+Dox** |
| **Nb GFP+ embryos/injected embryos** | **26/77** | **62/76** |
| Nb NANOG+ embryos/GFP+ embryos | 0/26 | 0/26 |
| Nb SOX2+ embryos/GFP+ embryos | 13/26 | 3/25 |
| Nb GATA6+ embryos/GFP+ embryos | 25/25 | 19/25 |
| **Total** | **143** | **158** |
| **Total injected embryos** | **301** | |

| | Rabbit host | |
|---|---|---|
| | E6 | E9 |
| Mouse ESCs | | |
| **Nb GFP+ embryos/injected embryos** | **9/9** | **12/20** |
| Nb SOX2+ embryos/GFP+ embryos | 6/6 | |
| Nb SOX17+ embryos/GFP+ embryos | 3/3 | |
| Rhesus TL2i | | |
| **Nb GFP+ embryos/injected embryos** | | **0/16** |
| **Total** | **18** | **36** |
| **Total injected embryos** | **54** | |

| | Cynomolgus host |
|---|---|
| | E7 |
| Rhesus TL2i | |
| **Nb GFP+ embryos/injected embryos** | **2/7** |
| Nb NANOG+ embryos/GFP+ embryos | 0/2 |
| hiPSCs_TL2i | |
| **Nb GFP+ embryos/injected embryos** | **2/7** |
| Nb NANOG+ embryos/GFP+ embryos | 0/2 |
| hiPSCs_t2iLGoY | |
| **Nb GFP+ embryos/injected embryos** | **2/15** |
| Nb NANOG+ embryos/GFP+ embryos | 1/2 |
| Mouse ESCs | |
| **Nb GFP+ embryos/injected embryos** | **3/7** |
| Nb NANOG+ embryos/GFP+ embryos | 1/1 |
| Nb SOX2+ embryos/GFP+ embryos | 2/2 |
| **Total injected embryos** | **36** |

Total embryos injected in this study : 3961

**Table S3: Summary of culture media used in this study**

| | E-NHSM | NHSM-v | 4i/L/b | TL2i | t2iLGöY | LCDM (EPS) | TL-CDK8/19i | 5i/L/A |
|---|---|---|---|---|---|---|---|---|
| **Media** | | | | | | | | |
| Neurobasal:DMEMF12 (1:1) | ✓ | | | | ✓ | ✓ | | ✓ |
| KO-DMEM | | ✓ | ✓ | ✓ | | | ✓ | |
| N2B27 | ✓ | | | | ✓ | ✓ | | ✓ |
| KOSR | 0.5% | 20% | 20% | 20% | | | 20% | 0.5% |
| **Transgene** | | | | | | | | |
| STAT3-ERT2 | | | | ✓ | | | ✓ | |
| **Growth factors** | | | | | | | | |
| LIF | 20ng/ml | 10ng/ml | 10ng/ml | house-made | 10ng/ml | 10ng/ml | house-made | 10ng/ml |
| Activin A | 20ng/ml | | | | | | | 20ng/ml |
| bFGF | | 8ng/ml | 4ng/ml | | | | | |
| TGFb1 | | 1ng/ml | | | | | | |
| **Small molecules** | | | | | | | | |
| BIRB796 | 2μM | | | | | | | |
| CGP77675 | 1μM | | | | | | | |
| CHIR99021 | 1.5μM | 3μM | 3μM | 3μM | | 1μM | | |
| PD0325901 | 1μM | 1μM | 0.5μM | 1μM | 1μM | | | 1μM |
| SP600125 | 5μM | 10μM | 10μM | | | | | |
| Gö6983 | 2μM | | | | 1μM | | | |
| SB202190 | | 5μM | | | | | | |
| SB203580 | | 10μM | 10μM | | | | | |
| (S)-(+)-Dimethindene maleate | | | | | | 2μM | | |
| Minocycline hydrochloride | | | | | | 2μM | | |
| IWR1 | 5μM | | | | | 0.75μM | | |
| XAV939 | | | | | 2μM | | | |
| CDK8/19i | | | | | | | 1.1μM | |
| IM-12 | | | | | | | | 1μM |
| SB590885 | | | | | | | | 0.5μM |
| WH-4-023 | | | | | | | | 1μM |

### Table S4: List of antibodies

| Target | Supplier | Reference | Host species | Dilution for cells | Dilution for pre-implantation embryo | Dilution for cryosection | Dilution for immunoblotting |
|---|---|---|---|---|---|---|---|
| GFP | Thermo Fischer | A10262 | Chicken | 1:300 | 1:300 | 1:300 | NA |
| mCherry | Thermo Fischer | M11217 | Rat | 1:300 | 1:300 | NA | NA |
| OCT4 | Santa Cruz | sc-9081 | Rabbit | 1:300 | 1:300 | 1:300 | NA |
| NANOG | R&D Systems | AF1997 | Goat | 1:100 | 1:100 | 1:100 | NA |
| NANOG | R&D Systems | AF2729 | Goat | NA | 1:100 | 1:100 | NA |
| SOX2 | R&D Systems | AF2018 | Goat | 1:100 | 1:100 | 1:100 | NA |
| SOX17 | R&D Systems | AF1924 | Goat | 1:100 | 1:100 | NA | NA |
| GATA6 | R&D Systems | AF1700 | Goat | 1:100 | 1:100 | NA | NA |
| SUSD2-PE | BioLegend | 327406 | Mouse | 1:100 | NA | NA | NA |
| KLF4 | Tebu-Bio | 09-0021 | Mouse | 1:100 | NA | NA | NA |
| TFE3 | Thermo Fischer | PA5-21615 | Rabbit | 1:250 | NA | NA | NA |
| KLF17 | Sigma-Aldrich | HPA024629 | Rabbit | 1:100 | NA | NA | NA |
| HuN | Sigma-Aldrich | MAB1281 | Mouse | NA | 1:100 | NA | NA |
| TuJ1 | Sigma-Aldrich | T2200 | Rabbit | NA | NA | 1:100 | NA |
| H3K27me3 | Cell Signalling | 9733 | Rabbit | 1:1000 | NA | NA | 1/1000 |
| TFAP2C | R&D Systems | AF5059 | Goat | 1:100 | NA | NA | 1/1000 |
| TFCP2L1 | R&D Systems | AF5726 | Goat | 1:100 | NA | NA | 1/1000 |
| CDK2 | Cell Signalling | E8J9T | Rabbit | NA | NA | NA | 1/1000 |
| CDK4 | Abcam | Ab199728 | Rabbit | NA | NA | NA | 1/1000 |
| CYCLIN E1 | Cell Signalling | 20808 | Rabbit | NA | NA | NA | 1/1000 |
| CYCLIN A2 | Thermo Fischer | MA1-154 | Mouse | NA | NA | NA | 1/1000 |
| CYCLIN D1 | Cell Signalling | 55506 | Rabbit | NA | NA | NA | 1/1000 |
| CYCLIN D2 | Cell Signalling | 3741 | Rabbit | NA | NA | NA | 1/1000 |
| CYCLIN D3 | Cell Signalling | 2936 | Mouse | NA | NA | NA | 1/1000 |
| p27kip1 | Cell Signalling | 3688 | Rabbit | NA | NA | NA | 1/1000 |
| ACTINB | Sigma-Aldrich | A3854 | Mouse | NA | NA | NA | 1/10000 |
